# The genome assembly of *Sphaerospora molnari* provides novel insights into the rapid evolution and diversification of unique lineage-specific gene groups in myxozoan parasites

**DOI:** 10.64898/2026.02.11.705288

**Authors:** Anush Kosakyan, Monika Wiśniewska, Enrique Riera-Ferrer, Gema Alama-Bermejo, Ivan Fiala, Anežka Karlíková, Ron P. Dirks, Martin Koliskó, Astrid S. Holzer

## Abstract

**Background:** Myxozoans are ancient cnidarian parasites with highly derived genomes characterized by an extremely accelerated rate of nucleotide substitution, abundant orphan and lineage-specific genes without further characterization. Genome data are limited to two out of the four main evolutionary lineages, assemblies are highly fragmented and often show significant levels of host contamination.

**Results:** We present a near chromosome-scale myxozoan genome, based on Oxford Nanopore-reads of *Sphaerospora molnari,* a member of the previously uncharacterized group of blood-feeding myxozoans, thereby addressing a key gap in the subphylum’s phylogeny. The haploid genome assembly spans 40.17 Mb in 40 contigs, harbors 14,957 genes and shows the smallest mitogenome of myxozoans (14,015 bp). Gene gain/gene loss analyses showed that myxozoans have small ancestral gene repertoires and show highly lineage-specific genome compositions. Using comparative analyses focusing on identifying unique but diversified gene compartments in *S. molnari,* we discovered taxonomically restricted protist genes related to red blood cell attachment (*Plasmodium* ETRAMPs) and surface protein variation (*Plasmodium* variant surface antigen families RIFIN and STEVOR, as well as Metamonada variant-specific surface proteins, VSPs), raising questions about their origins and evolution. A genomic trait shared between several myxozoans is the significant expansion of DNA transposable elements belonging to the mutator-like elements (MULEs), and while the simple copy-paste mechanism of these transposases may suggest frequent uncontrolled mutation, we demonstrate domestication of MULEs into transcription factors. Analyses of the gene fragments of chimeric MULEs (pack-MULES) of *S. molnari* show that these coincide with highly diversified gene groups in this lineage, including alien genes, suggesting MULEs as a driving force for gene evolution in myxozoans.

**Conclusions:** Parasitic lifestyle shifts drive exceptionally rapid genome evolution in myxozoans, primarily through nonadaptive mutation and gene transfer via MULEs, with adaptive refinement through MULE domestication and selection. In *S. molnari*, these processes underpin unique erythrocyte exploitation and immune evasion strategies essential for survival in the hosts’ bloodstream.

## Background

Myxozoans are an exceptionally successful, diverse, and in many aspects enigmatic group of endoparasitic organisms with an obligatory two-host life cycle, alternating between aquatic vertebrates (mainly fish) and invertebrates (annelids and bryozoans). Multicellular spores serve as transmission stages in both, marine and freshwater aquatic habitats. Spores contain extremely simplified nematocysts (polar capsules), used for host attachment; these structures constitute the only clearly recognizable feature linking myxozoans to the phylum Cnidaria, to which they belong. Together with extensive reduction of cell layers, body polarity and organs to simple, pluricellular stages, myxozoans have undergone substantial gene losses resulting in miniaturized genomes.

Genome size in cnidarians averages ∼470 Mb, ‘giants’ can be found amongst Siphonophores, estimated at 0.7-2.3 Gb [1], while myxozoans possess the smallest genomes within the phylum, at only 12.9-185 Mb [2–5]The genome of the myxozoan *T. bryosalmonae* (12.9 Mb) is likely the smallest metazoan genome known to date, followed by that of the orthonectid *Intoshia variabili* (15.3 Mb; [6]), a parasite of marine invertebrates. Genome reduction through gene loss and compaction is common to many parasite groups [7]. However, parasites must adapt to specific hosts, niches and nutrients and hence ‘make’ different choices in terms of which genes to keep and which to lose. Some gene families will have to expand to improve the parasite’s ability to infect and survive within a host.

Myxozoans exhibit particularly extensive gene loss, including genes considered hallmarks of metazoan development [8], such as loss of the majority of core apoptotic domains [9], key oxygen-sensing and homeostasis genes [10], and in one species - the only known metazoan case – genes involved in aerobic respiration and mitochondrial genome replication [11]. Although myxozoan evolution has overall been conceptualized as a process of genetic loss and simplification, the genome of *Myxobolus honghuensis*, a tissue-dwelling myxozoan parasite, reveals a mosaic architecture of conserved, depleted, but also enhanced genes and pathways [4]. While the number of expanded and contracted gene families is similar, the expanded families contain ∼23-fold more genes, suggesting large-scale gene duplication. Moreover, a disproportionally large percentage of genes in myxozoan genomes code for unknown proteins, some highly abundant in their datasets [12]. Such genes can confer novel biological functions and represent an important source of evolutionary innovation, enabling physiological adaptation and ecological transitions.

Over the past decade we have established *Sphaerospora molnari* as a laboratory model for myxozoan research. *S. molnari,* a parasite of common carp in Europe, belongs to one of the four major phylogenetic clades of myxozoans, the *Sphaerospora* sensu stricto clade [13]. A lineage-specific trait identified only in this clade is the presence of extracellular stages that circulate and proliferate in the blood stream of their fish hosts (see [14] and references therein). We discovered that *S. molnari* attaches to and feeds on red blood cells, inducing hemolytic anemia [15]. Although blood is a nutrient-rich resource for parasites, immune cells, amongst which nucleated erythrocytes play important role, especially in teleosts [16,17], create a hostile environment. Parasites therefore require effective defense, immune evasion, or immune-modulatory strategies to avoid recognition and elimination. During its intrapiscine life cycle, *S. molnari* also localizes in the liver, while spore-formation takes place in the gills [18], indicating its capacity for target site recognition and site-specific development. The unusual developmental pattern, including proliferative blood stages, distinguishes *S. molnari* from all myxozoans for which genomic data exist to date, as these belong to distantly related phylogenetic clades of predominantly tissue-dwelling or bile-inhabiting species. The possibility to isolate the parasite at 99.9% purity from the host [19] combined with advanced Oxford Nanopore technology sequencing provided a unique opportunity for producing a high-quality myxozoan genome for comparative analyses.

Myxozoans evolved 650 mya and are some of the oldest metazoans populating our planet; they have co-evolved with and adapted to their fish hosts since their acquisition more than 400 my [20]. To elucidate the genetic basis underlying their evolutionary success and extensive diversification, we aimed at identifying genes unique to myxozoans but shared across major lineages. In parallel, we sought to characterize the unique adaptive gene repertoire of sphaerosporids that enables their survival and proliferation within the hostile environment of the fish bloodstream.

## Results and Discussion

### A near-chromosome-scale genome assembly for *S. molnari*

We sequenced and assembled the genome of the myxozoan *S. molnari* using high-coverage long nanopore reads (8.2 million reads before filtering, 35x coverage after filtering) assisted with Illumina paired-end sequences (293 million paired-end reads before filtering). Using 99.9% pure, DEAE-column isolated *S. molnari* blood stages as a source [19], filtration of the obtained sequences yet resulted in a recovery of only 61.7% parasite data alongside 38.3% fish host contamination. This contamination is likely caused by the parasite feeding on host red blood cells and the transfer of host materials, including DNA to the parasites [15], and by the massive (approx. 39-fold) difference in genome size between host and parasite (host genome [21]). The haploid post-filtration assembly of *S. molnari* spans 40.17 Mb and contains 40 contigs, many of which are near-chromosome size (smallest contig is ∼46 kb; N50=2.8 Mb; genome assembly accession number GCA_046867295.1, Bioproject PRJNA1193866). *S. molnari* has an average genome size when compared to other myxozoans for which genome assemblies are currently available on GenBank (17 taxa, 20 assemblies). Two assemblies are considerably larger, that of *Thelohanellus kitauei* (150.3 Mb), the first myxozoan genome published, and that of *Myxobolus honghuensis* (161 Mb; [4]). The larger assembly size of these two species may in part be a result of the sequencing technology used at the time (454, Illumina), however these assemblies also contain considerably more predicted genes (16,638 and 15,433) than all other species (5,316-9,716). The *de novo* assembled genome of *S. molnari* is close to complete and continuous, as 98.94% of transcripts mapped to the genome along their full length with an identity ≥ 90%. Genome structural annotation predicted 14,957 genes, the highest number of genes in relation to assembly size in myxozoans. Genome completeness of *S. molnari* was 64% (163/255 genes; complete 163 (single copy 158, duplicated 5), fragmented 27, missing 65) based on the Eukaryota Odb10 single-copy ortholog dataset, which represents conserved genes encoding core biological functions [22]. This score is lower than typically observed in free-living cnidarians. The strongly accelerated rate of molecular evolution in myxozoans, possibly the fastest known among eukaryotes [20] may hinder the detection of eukaryotic core genes [11]. However, reductions may also have occurred in parallel with overall genome size contraction. *S. molnari* possesses the largest complement of eukaryotic core genes reported in Myxozoa (163; this study), followed by *Ceratomyxa* sp., Sample 70 (123), but no more than six orthologs appear to be shared among taxa [23], likely due to lineage-specific losses. The relatively large number of eukaryotic core genes in *S. molnari* may be reasoned by its relatively basal phylogenetic position (see below).

Eight of 19 myxozoan genome assemblies available on GenBank were published in the last 24 months, and, in contrast to previous submissions, these datasets were produced by long-read sequencing technologies (Oxford Nanopore, PacBio). The combination of PacBio and Arima technologies recently resulted in the chromosome-scale assembly of *Kudoa neothunni* as a byproduct of sequencing endeavors originally aimed at the host genome [24]. The importance of novel sequencing technologies cannot be underestimated in this parasite group where genome organization, the repeat landscape and patterns of rearrangement are still largely unknown. The *S. molnari* genome assembly combines high quality with near-chromosome-scale contiguity and, most importantly, fills a crucial gap in the phylogenetic landscape of the subphylum Myxozoa.

In addition to long-read sequences we obtained physical data on nuclear chromosomes in arrested metaphase. As characteristic for myxozoans, *S. molnari* proliferative blood stages represent cell-in-cell complexes with at least one primary cell holding one secondary cell, up to compositions of many secondary cells holding tertiary cells, all within one primary cell (**Figure 1a**). We found primary cells were non-dividing, but we were able to visualize 7 chromosomes (2n=14, **Figure 1b**) in secondary cells undergoing mitotic division (**Figure 1c**). Chromosomes were found to be very small with approx. 0.6µm length, 4 to 10 times smaller than the average chromosome size in free-living cnidarians (e.g. [25][26]). The high chromosome number observed in *S. molnari* stands in contrast to the commonly reported karyotype of myxozoans, which typically consists of four chromosome pairs (2n = 8) in reports from the genera *Ceratomyxa, Zschokkella, Myxidium, Henneguya, Myxobolus, Sphaeromyxa* [27–30]. Lower counts of three chromosome pairs (2n = 6) (2 reports; [24,31]) and higher counts of six chromosomes (2n=12) (3 reports; [32–34]) seem to be exceptional. Fluorescent *in situ* hybridisation of 18S rDNA in *S. molnari* demonstrated its localisation on a single chromosome pair (2 chromosomes; **Figure 1d**). In *S. molnari*, FISH visualization corresponds with genome sequence data where rDNA is located at the ends of contig_14 (504 kb) and contig_30 (1.34Mb). The inability to bridge these contigs is likely due to arrays of tandemly repeated rDNA units within this genomic region. Such rDNA clusters are consistent with the high rRNA gene copy numbers previously reported for other myxozoans, including >100 copies in *C. shasta* [35] and >1000 in *Ceratomyxa puntazzi* [36].

**Figure 1.**
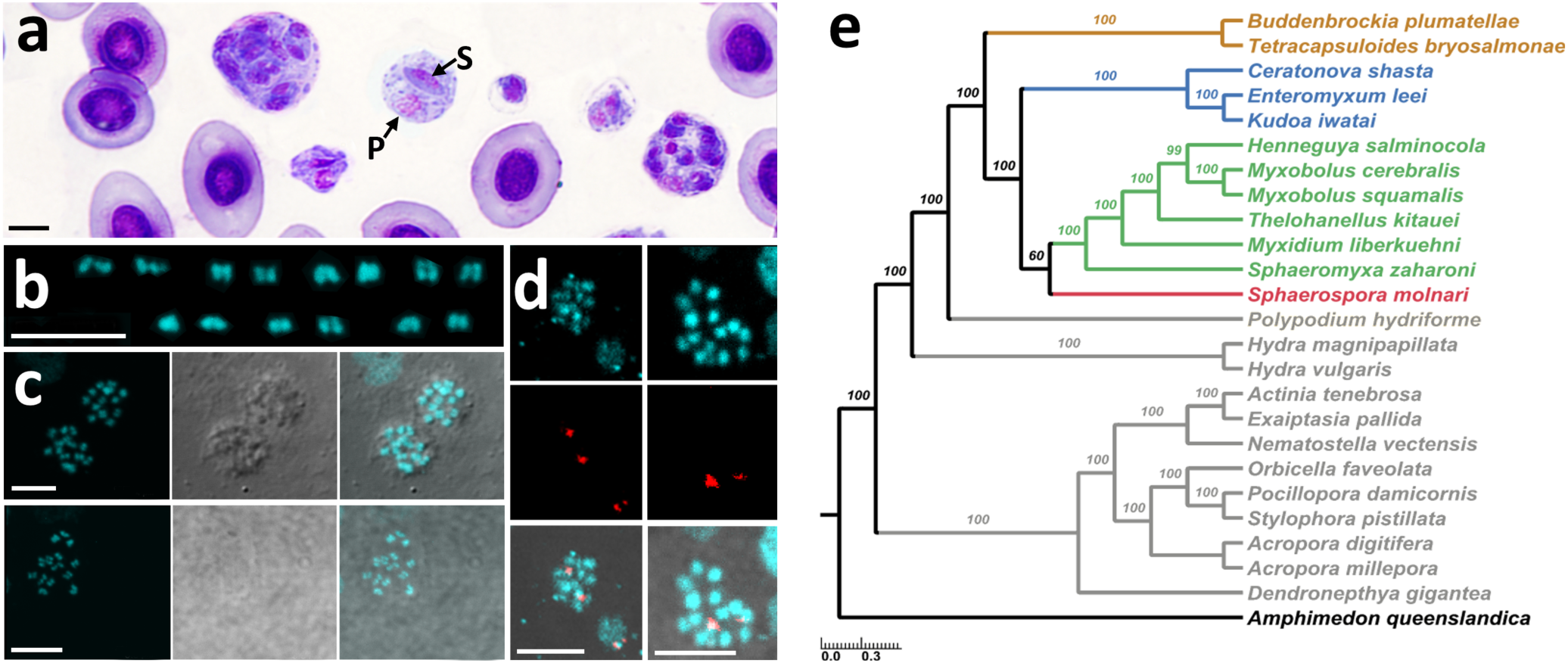
*S. molnari* morphology, chromosome data and phylogenomics: **a.** Blood smear of common carp showing *S. molnari* multicellular blood stages composed of 2-13 small cells (and nuclei) in amongst red blood cells (cells with single large nuclei), showing typical myxozoan cell-in-cell structure with P=primary cell, S=secondary cell and some larger stages with tertiary cells (not indicated), Diff-Quik staining. **b.** Chromosome map of *S. molnari* showing 7 pairs of small chromosomes in arrested metaphase (DAPI-stained). **c.** Secondary cell nuclei of *S. molnari* with chromosomes in arrested metaphase (left=DAPI stain, middle=DIC light microscopy, right=combination of the two: upper set=two nuclei, lower set=single nucleus). **d.** ISH using an 18S rDNA probe (Cy3-streptavidin, red fluorescence) binding to two loci in diploid *S. molnari* chromosomes (DAPI stained); (top row=DAPI, 2^nd^ row=Cy3-streptavidin, bottom row=combination). Bars in **a-d**=4 µm. **e.** Phylogenomic position of selected cnidarians based on Maximum Likelihood analyses of 77 ribosomal protein-coding genes (12,809 aa). Myxozoans (colorful lineages) are reconciliated as a monophyletic group, sister to the parasitic Polypoidididae (*Polypodium hydriforme*). Within the Myxozoa, malacosporeans (myxozoans infecting bryozoans; brown) cluster basal to the polychaete-infecting lineage (blue) and the oligochaete-infecting lineage (green) which includes *S. molnari* (red) in a basal, weakly supported position. Free-living cnidarians are shown in grey. Outgroup: *Amphimedon queenslandica* (Porifera; black); values at bifurcations indicate bootstrap support estimated from 100 PMSF replicates.

We did not achieve binding of the universal FISH telomeric probe sequence [TTAGGG]n to *S. molnari* chromosomes, despite the generally conserved nature of this sequence in basal metazoans [37]. However, we were also unable to identify TTAGGG repeats bioinformatically, and we believe that telomeric sequences of *S. molnari* and other myxozoans vary from the general metazoan repeat sequence. In *S. molnari* we were able to identify a consensus hexamer repeat of [CYAACY]n at the end of multiple long contigs. We believe that telomere repeats and subtelomeric regions require further investigation in myxozoans due to the contingency genes located in this area, an organization which show parallels to e.g. *Plasmodium* [38].

### Phylogenomics and a single-circle mt genome identify *S. molnari* as a basal annelid-infecting lineage

Myxozoan phylogenies have been based predominantly on 18S rDNA sequences [39,40], and it has been suggested that the main four major lineages likely emerged in different invertebrate host groups [20]. The invertebrate host of sphaerosporids is only known for one species [41], and remains without molecular proof of identity of the two morphologically distinct spore stages in its life cycle. The unstable positioning of the three more recent clades, sphaerosporids, oligochaete-infecting and polychaete-infecting myxozoans to each other [13] may be caused by missing sequences that would link ancient host acquisition events in co-evolving parasites and fish hosts [20]. In the present study, the analysis of a dataset composed of 77 protein-coding genes resolved myxozoans as a monophyletic lineage with 100% ML bootstrap support, and as a sister group of *Polypodium hydriforme* (Polypodidae:Cnidaria) (**Figure 1e**), similar to previous phylogenomic studies [42],[43]. The overall topology in our tree shows malacosporeans (myxozoans using bryozoan hosts) as the most basal group. However, in contrast to previous analyses based on 18S rDNA, in the present study, *S. molnari* clustered basal to oligochaete-infecting (predominantly freshwater) species and sister to polychaete-infecting (predominantly marine) species (**Figure 1e**). Its basal and weakly supported topology (boostrap support=60) may indicate that *S. molnari* represents a myxozoan lineage that originated in a different annelid stem group, as previously suggested by Patra et al. [44]. To improve our understanding of sphaerosporid evolution, additional genome assemblies from other members of the group are essential. Data from more basal marine species will be critical for addressing current phylogenetic ambiguities.

The ancestral myxozoan mt genome is believed to be a single chromosome of circular morphology [45], while some derived myxozoan mt genomes have been shown to possess mt plasmids or mt minicircles, which are likely the result of recombination between repeated elements [45,46]. Potentially further confirming its basal phylogenetic position, the *S. molnari* mt genome is encoded on a single circle with a genome length of 14,015 bp (GenBank accession number PQ893892; **Figure 2a**). It is the smallest mt genome reported in myxozoans (**Figure 2b**), with around 60% representing non-coding regions, and an overall GC-content of 27.2%. Metazoan mt genomes generally contain 13 protein-coding genes, and so do those of most free-living cnidarians [47]. Myxozoan mt genomes are reduced and usually contain only seven canonical genes (**Figure 2b**): five protein-coding genes, i.e. cytochrome b (*cob*), the cytochrome oxidase genes *cox1* and *cox2*, NADH dehydrogenase subunits 1 and 5 (*nad1* and *nad5*), and two rRNA genes (*rnl* and *rns*). *Myxobolus squamalis* lacks *cox1*, *cox2* and *cob*, and *Henneguya salminicola* lacks the mitochondrial genome entirely and is unable to perform mitochondrial respiration [5,11,45]; and those cited herein). In *S. molnari*, we only identified five of the seven canonical mt genes: *cox1*, *cox2*, *cob*, and the two rRNA genes (*rnl* and *rns*) (**Figure 2a**). Neither the two canonical *nad1* and *nad5* nor other NADH dehydrogenase genes, previously found solely in histozoic intracellular myxozoans (*Kudoa* spp., [48]) were detected (**Figure 2b**). Alignment of RNA reads to the mitochondrial genome confirmed the *rnl* locus, but insufficient read-length coverage over the *rns* region (**Supplementary figure 1**) resulted in a tentative annotation of this gene based on MITOS2 output. In *S. molnari,* we found 2 unidentified ORFs (>100aa) that were observed to be transcribed and to contain putative transmembrane domains, suggesting an involvement in the electron transport chain. Similarly, smaller *Kudoa* spp. mt genomes (15,880-19,350 bp) have an average of 3 unidentified ORFs [48]. These appear to align with unidentified ORFs found in *S. molnari*. Concatemers of mtDNA detected in *S. molnari* long reads suggests the existence of an intermediate molecule of a “rolling-circle”-like mechanism of mtDNA replication, containing one complete copy and two partial copies of mtDNA. Interestingly, in yeast and humans, mt DNA concatemers are often observed in conditions of oxidative stress, with elevated reactive oxygen species (ROS) levels associated to processes such as cellular proliferation and apoptosis [49,50].

**Figure 2.**
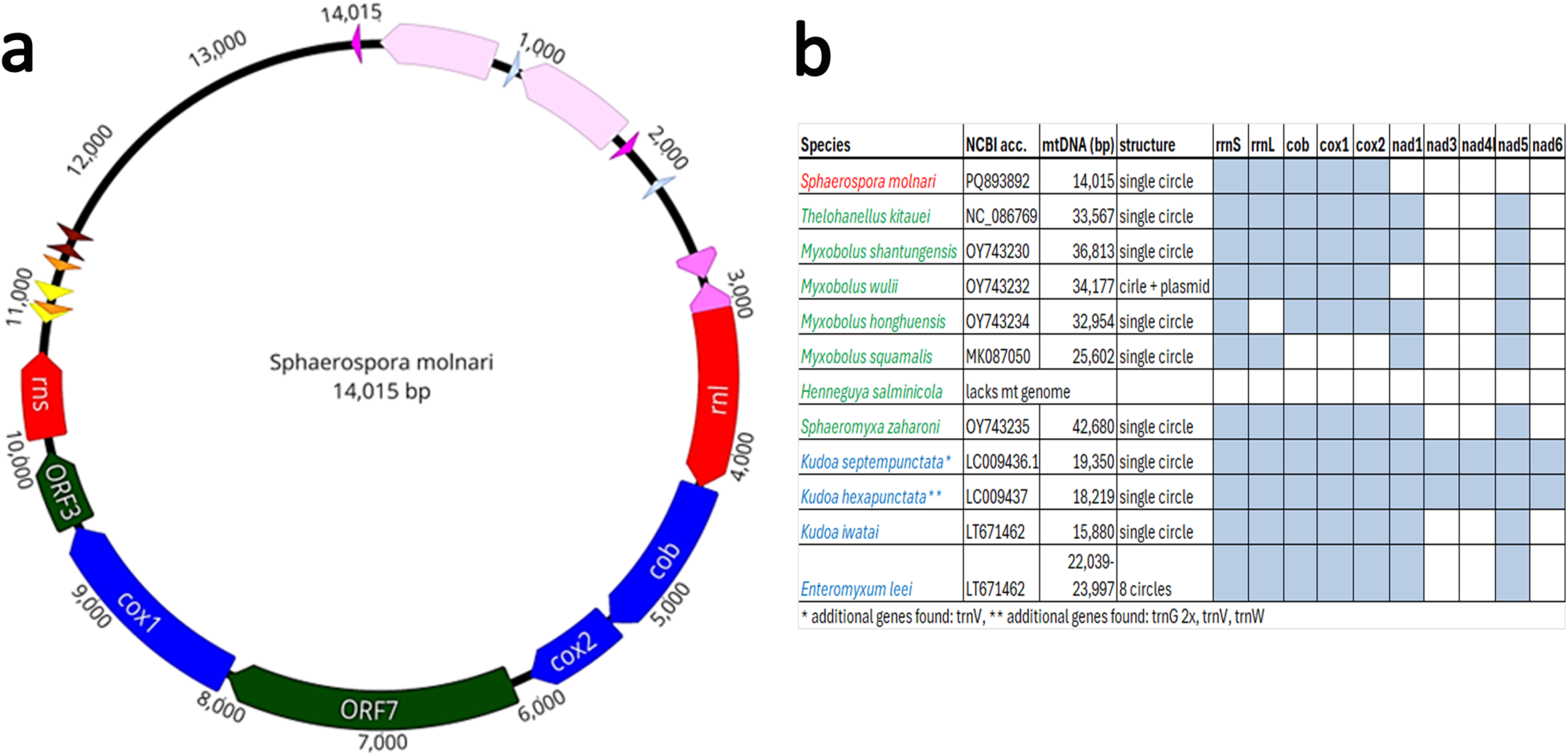
The mitogenome of *S. molnari* in comparison with that of other myxozoans. **a.** *S. molnari* mt genome map (14,015bp). The three canonical mt protein coding genes (*cox1, cox2* and *cob*) are in dark blue, the two rRNA subunits (*rns* and *rnl*) are in red, unknown ORFs in green. All other arrows indicate repeat elements. Repeat elements with the same sequence have the same color. Data produced in MITOS2. **b.** mt genome features including genome size, gene content and structure in different myxozoans, in comparison (lineage-specific color coding of taxa as in **Figure 1b**).

### *S. molnari* has the most metabolically complete myxozoan genome featuring a sugar-based energy acquisition strategy

Comparative analyses of myxozoan KEGG metabolic pathways reveal extensive metabolic reduction across Myxozoa, consistent with their parasitic lifestyle. *S. molnari* emerges as the metabolically most complete and versatile taxon in the dataset analysed. It retains comparatively intact carbohydrate, lipid, energy, and nucleotide metabolism pathways that are highly reduced or absent in other myxozoans, likely reflecting its systemic, blood-borne lifestyle, access to host nutrients, and its phylogenetically basal position within the group. Notably, *S. molnari* preserves a near-complete central carbohydrate metabolism, including glycolysis/gluconeogenesis (M00001), the citrate cycle (M00009), the pentose phosphate pathway (M00004–M00007), and pyruvate metabolism/oxidation (M00307), enabling aerobic energy production, redox balance, and synthesis of glucose-6-phosphate as a key metabolic hub. These genomic features are consistent with experimental observations that exogenous glucose is required for successful *in vitro* culture of *S. molnari* [19].

In contrast to other myxozoans, which show fragmented or incomplete nucleotide and lipid biosynthetic pathways, *S. molnari* retains complete *de novo* purine and pyrimidine synthesis pathways, whereas other taxa lack multiple modules or exhibit disrupted interconversion steps. Although myxozoans generally lack canonical *de novo* fatty acid biosynthesis and elongation pathways, indicating a strong reliance on host-derived lipids, *S. molnari* exhibits broader lipid metabolic capacity, including glycerophospholipid metabolism (M00032, M00033), sphingolipid metabolism (M00067–M00071), and triacylglycerol metabolism (M00061). This expanded lipid repertoire is consistent with the presence of phospholipases and cholesterol acyltransferases, supporting fatty acid degradation, energy storage, membrane biogenesis, and lipid-mediated signaling. Furthermore, *S. molnari* uniquely retains complete oxidative phosphorylation and electron transport chain modules (M00142–M00158), indicating intact mitochondrial function and active aerobic respiration, whereas other myxozoans such as *Ceratonova shasta*, *Thelohanellus kitauei*, *Henneguya* and *Myxobolus* spp. show severe pathway fragmentation. While amino acid biosynthesis is broadly reduced across Myxozoa, *S. molnari* retains select pathways, including polyamine biosynthesis (M00855), proline biosynthesis (M00015), and cysteine and glutathione metabolism (M00338, M00415), which are associated with cellular homeostasis, oxidative stress defense, and host–parasite metabolic interactions [51].

### *S. molnari* combines an ancestral gene repertoire with a highly lineage-specific genome composition

The OrthoFinder comparative analysis, which included genomes and transcriptomes from free-living cnidarians (nine species), eight myxozoans, and *Polypodium hydriforme*, recovered 32,711 orthogroups (OGs), of which 855 are conserved across all taxa. Free-living cnidarians possess 8,156–12,372 OGs, reflecting their more complex gene repertoires, whereas myxozoans show substantial reductions, retaining only 3,483–6,141 OGs (average contraction of ∼55%). The *S. molnari* genome has 3,658 OGs represented by 9,139 genes (excluding 5,818 unassigned genes). Of these, 3,195 OGs (5,422 genes) are shared with at least one other myxozoan species (**Figure 3a**). Gene repertoires varied markedly among myxozoans, with only 138 OGs shared across all eight taxa included (**Figure 3a**), underscoring extensive lineage-specific gene loss and gain. The *S. molnari* genome demonstrates substantial ancestral cnidarian gene content but also possesses a notable set of 550 species-specific OGs (3,973 genes) (**Figure 3a**) resulting in multiple unique clusters (>77%) in its protein architecture network (**Figure 3b**). Only *T. kitauei* and *S. zaharoni* have a higher share of lineage-specific OGs (831 and 826 OGs) (**Figure 3a**).

**Figure 3.**
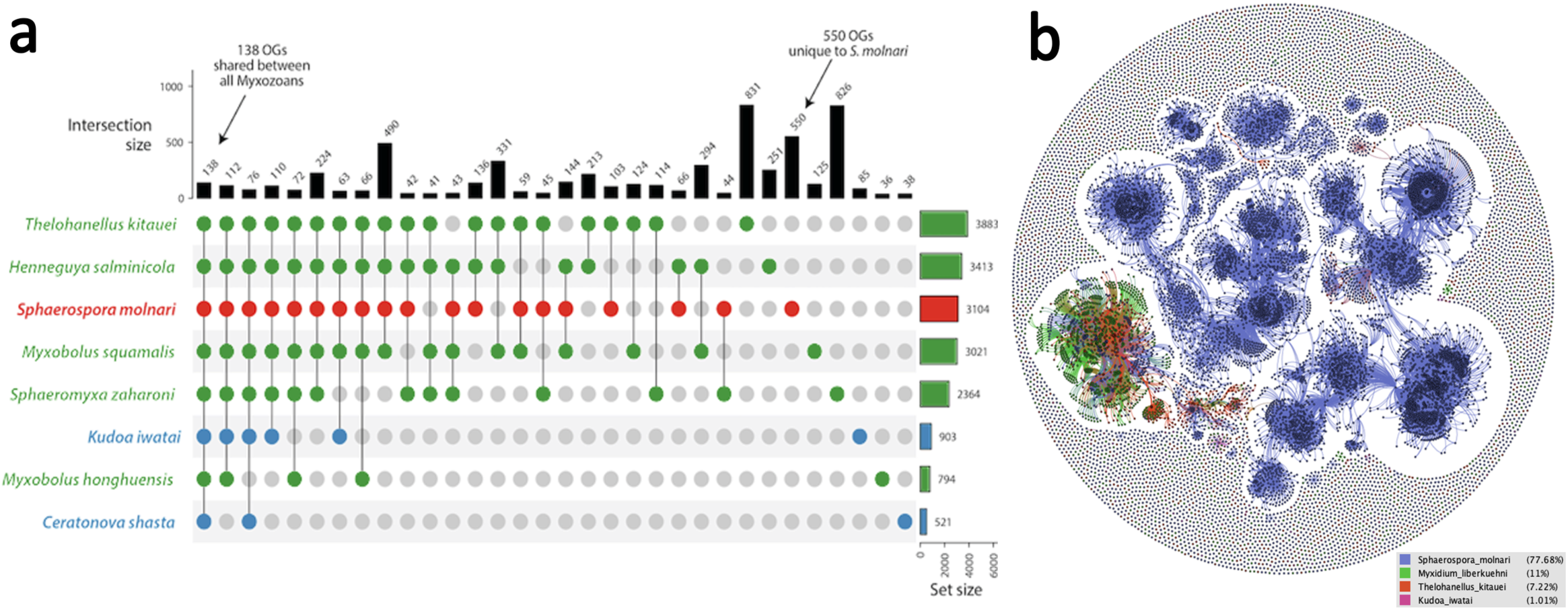
Lineage-specific gene innovation in the Myxozoa. **a.** UpSetPlot (Python) depicting orthogroups (OG) shared among myxozoan taxa: Only 138 OGs are shared between all 8 myxozoan taxa selected for analyses. *S. molnari*, *T. kitauei* and *S. zaharoni* show the highest number of unique orthogroups (550-831). **b.** Orthology-based comparative protein network (Orthofinder, Cytoscape) with *S. molnari*-unique protein clusters in blue. Clusters shared with other species in different color, indicated for species where the overlap is >1.

This indicates rapid acquisition of genomic novelty in these species, which appears more pronounced in myxozoans from the oligochaete-infecting (freshwater) lineages, than in those from the polychaete-infecting (marine) lineage (**Figure 3a**). It is obvious that the Myxozoa show extensive gene content variations and that functional differences among recently gained and lost genes likely represent the basis of distinct host exploitation strategies and evolutionary trajectories.

Evidence of the relevance not only of newly acquired but also of re-destined genes in myxozoan genomes can be found in their duplication rate. In *S. molnari,* 3,306 genes/domains were ≥5x duplicated; a substantial amount, 2,912 of these, are unique to this species. Amongst the 41% annotated genes, a very large percentage of genes/domains is related to DNA transposition via TEs, functions as transcription factors, or represent membrane proteins with specific functions.

### Host-parasite interaction and immune regulating genes specific to the Myxozoa

Several genes involved in host-parasite interaction and immune regulation are duplicated in the majority of myxozoans and hence constitute essential adaptive strategies for successfully inhabiting fish hosts. An important group are **arginine repressors**, duplicated e.g. in *Myxidium lieberkühni, T. kitauei,* and especially in *S. molnari*. These repressors regulate the transcription of arginine metabolic pathways. Arginine plays a central role in immune regulation by activating cytotoxic macrophages, natural killer cells, and T cells, and by modulating chemokine signaling. Macrophage function during immune responses is largely determined by arginine metabolism [52,53]. Pro-inflammatory M1 macrophages express nitric oxide synthase, converting arginine into nitric oxide (NO), a potent cytotoxic molecule, whereas anti-inflammatory M2 macrophages express arginase, which hydrolyzes arginine to ornithine, promoting wound healing and parasite proliferation via the synthesis of polyamines [52,54]. Several pathogens actively reduce host arginine availability. For example, anaerobic prokaryotes and metamonad protists such as *Giardia intestinalis* use the arginine deiminase pathway to consume arginine, sometimes as a major energy source [55]. In *G. intestinalis*, arginine depletion has been linked to reduced epithelial turnover, promoting a stable niche for long-term infection [56]. While none of the enzymes of the arginine deiminase pathway were detected in myxozoans, we identified multiple gene duplications in arginine repressors. We hypothesize that myxozoans modulate host immune responses by regulating arginine availability, thereby promoting immune tolerance and long-term persistence at low parasite densities, despite the development of specific host immunity reported for several species [18,57]. This strategy may be particularly important for blood-dwelling parasites, potentially explaining the further expansion of arginine repressor genes in *S. molnari*, where suppression of host immune effector mechanisms could prevent parasite elimination, as described in trypanosomes [58].

Another group of highly duplicated genes common to myxozoans are those involved **in signaling pathways involving mitogen-activated protein kinase (MAPK), phosphoinositide 3-kinases (PI3K), and transforming growth factor beta (TGF-β).** These pathways are generally involved in cell growth, proliferation, differentiation, migration and intracellular trafficking, apoptosis and homeostasis, as well as immune mechanisms and tolerance. In parasites they have been found important for disease development and parasite maintenance in the host. Different groups of intracellular parasites (i.e. *Leishmania, Apicomplexa*) use MAPK and PI3K pathways to evade apoptosis and persist and survive inside cells [59,60]. This may explain why myxozoans with long-term intracellular developmental stages (*Kudoa* spp.) show higher duplication rates of the genes related to these pathways than species whose intrapiscine life cycle is dominated by an extracellular lifestyle. In *Schistosoma mansoni*, a novel parasite-encoded TGF-β superfamily member controls embryo development and host-parasite interaction [61], while in another helminth parasite, *Heligmosomoides polygyrus* a TGF-β mimic binds to the TGF-β receptors and stimulates the differentiation of naïve T-cells into T regulatory cells [62]. Tregs are essential for maintaining peripheral tolerance and limiting chronic inflammatory processes. Interestingly, myxozoans also demonstrate gene duplication of **epidermal growth factor (EGF)** domains, found in the extracellular portion of transmembrane proteins or in secreted proteins engaged in protein-protein interaction. Several were annotated with an EggNOG function for sequestering (likely host-derived) TGF-β in the extracellular matrix thereby regulating the bioavailability of TGF-β [63].

### Lineage-specific genes in *S. molnari*: surface proteins for immune evasion and host cell attachment derived from parasitic protists?

Amongst the genes unique to *S. molnari* the most intriguing discovery are those coding for 3 different types of highly duplicated membrane proteins, that show high similarity to taxonomically restricted genes described in specific lineages of parasitic protists. Their presence likely explains how sphaerosporids manage to persist and proliferate in the blood where both cellular and humoral immune factors are highly effective.

We identified 18 **variant-specific surface proteins (VSPs)**-coding genes in the genome of *S. molnari*. VSPs constitute a family of cysteine rich proteins (> 10% cysteine), characterized by the presence of an N-terminal signal peptide (20-25 aa), a large extracellular ectodomain of > 5 CXC or CXXC motifs (300-700 aa), a single transmembrane (TM) domain, followed by fewer than 100 amino acids and a conserved C-terminal motif (5-15 aa) [64]. To date, VSPs have been reported exclusively from diplomonads, where pathogenic species harbor substantially larger VSP repertoires than free-living taxa, with up to 106 VSPs in *Giardia intestinalis* and 111 in *Spironucleus salmonicida* [65]. Differential expression of VSP genes is triggered by an epigenetic mechanism and results in antigenic variation, allowing prolonged persistence of *Giardia* within the host intestine [66]. Although VSP aa sequences are generally highly divergent, *S. molnari* VSPs display clear structural similarities to diplomonad VSPs, including conserved overall architecture and a lineage-characteristic C-terminal signature motif [KR]X[KR]XK[KR], suggesting homology. *S. salmonicida* and *Hexamita* sp. are fish parasites, raising the possibility that sphaerosporid VSPs were acquired via lateral gene transfer from diplomonad fish parasites. Regardless of their origin, this study represents the first report of VSPs outside the Diplomonada. Evidence of antigenic variation during parasite development has previously been documented in *S. molnari* [18] and a few other myxozoans [67,68] but the underlying molecular mechanisms remained unknown. In *S. molnari* antigenic variation is particularly relevant as an immune evasion strategy given that the parasite proliferates in the blood and is therefore exposed to multiple immune effector mechanisms, including antibody-mediated responses [18].

The **repetitive interspersed family/subtelomeric variable open reading frame family (RIFIN/STEVOR)** is a *Plasmodium*-restricted **variant surface antigen (VSA)** family (∼190 genes; [69]) with gene homologues in *S. molnari* and important gene expansion by duplication (27 genes). **STEVOR** is an erythrocyte-binding protein recognizing Glycophorin C on the RBC surface [70] and is likely essential for *S. molnari* attachment to RBCs. It was shown that STEVOR is clonally variant at the surface of malarial schizont stage parasites. Expression of different STEVOR variants on the surface of the infected red blood cell changes the parasite’s antigenic property [70]. Similarly, **RIFINs** encode a large family of *Plasmodium*-specific transmembrane proteins which are expressed on the surface of infected red blood cells. Type A RIFINs were shown to bind to inhibitory immune receptors to evade NK and B cell recognition [71] by binding to inhibitory immune receptor KIR2DL1 more strongly than KIR2DL1 binds to the human ligand, MHC class I [72]. Like VSPs, the selective expression of VSAs enables *P. falciparum* and potentially *S. molnari* to evade immune clearance in the blood [73].

The third family of membrane proteins unique to *S. molnari* and of particular interest to host-parasite interaction are those belonging to **early transcribed membrane proteins** (ETRAMPs; 10 genes). Also taxonomically restricted to *Plasmodium*, ETRAMPs are developmentally regulated and highly charged membrane proteins located in the parasitophorous vacuole membrane (PVM) of malaria parasites, directly at the parasite-host cell interface [74]. In contrast to *Plasmodium*, *S. molnari* develops extracellularly but actively attaches to RBCs [15]. Highly charged molecules such as ETRAMPs may be responsible for interaction with structural proteins on the RBCs surface to which *S. molnari* attaches tightly for transfer of host proteins to the parasite [75]. Alternatively, myxozoan ETRAMPs may act as membrane carriers of soluble proteins in the parasite cytoplasm for effective transport outside the parasite cells.

The number of taxonomically restricted genes from parasitic protists that can be found embedded in the *S. molnari* genome raises questions on their origin and acquisition. High sequence and structural similarity suggest acquisition by LGT rather than convergent evolution. Analyses of the transposon repeat landscape of *S. molnari* supports LGT as a potential mechanism for acquisition (see below).

### Essential repeats: Myxozoans evolved unique drivers for an unmatched rate of molecular evolution by expansion of DNA transposons belonging to the MULE family

Repetitive DNA sequences play critical roles in genome evolution by promoting genetic variation and influencing gene regulation. Although accurate repeat quantification of repetitive sequences can be challenging in assemblies based exclusively on Illumina short-read data, which frequently fail to resolve repetitive regions accurately [76], we observed no consistent difference in overall repeat content between myxozoans and their free-living cnidarian relatives. This pattern is consistent with observations reported for other parasitic lineages. Intriguingly, repeat landscapes in Myxozoa exhibit pronounced shifts in composition, with certain repeat classes being markedly reduced while others show substantial diversification (**Figure 4a**). Among the genes unique to myxozoans, transposable elements (TEs) and transcription factors (TFs) represent the most diversified functional categories, together accounting for approximately 28% of all annotated genes. TEs are highly dynamic repeat elements that are capable of reshaping genome architecture by promoting rearrangements that lead to gene creation or disruption, gene shuffling, and modulation of gene expression patterns [77]. A defining characteristic of TEs is their strong propensity to induce mutations, thereby contributing to genome plasticity and evolutionary innovation [78]. Within transposable elements (TEs), most retrotransposons (class I TEs), including SINEs, LINEs, and long terminal repeat (LTR) elements, are nearly absent in the Myxozoa, except *Gypsy/DIRS1*. In contrast, DNA transposons (class II TEs) are generally reduced but display lineage-specific expansions in several families, particularly the *hAT, pogo,* and Mutator-like transposable element (MULE) superfamilies (**Figure 4a**). *hAT* (hobo-Activator) and *pogo* (Tc1-IS630-Pogo) occupy on average 2.3% and 3.5% of myxozoan genomes vs 0.6% and 0.3% of the genomes in free-living taxa. As much as 8.5% of *S. molnari’*s genome represents Tc1-IS630-Pogo, in *Myxobolus rasmusseni* the percentage even amounts to 12.3%. In polychaete-infecting species both classified and unclassified families of DNA transposons are strongly reduced in number in contrast to oligochaete-infecting species (**Figure 5a**). The mechanism of transposase-mediated DNA cleavage and target joining depends on the relevant pathway [79]: while MULEs cut the first DNA strand, in myxozoans the *hAT* and *pogo* superfamily seem to be responsible for cleavage of the second DNA strand (highly diversified class II TEs), while *piggyBac* (alternative pathway) is not.

**Figure 4.**
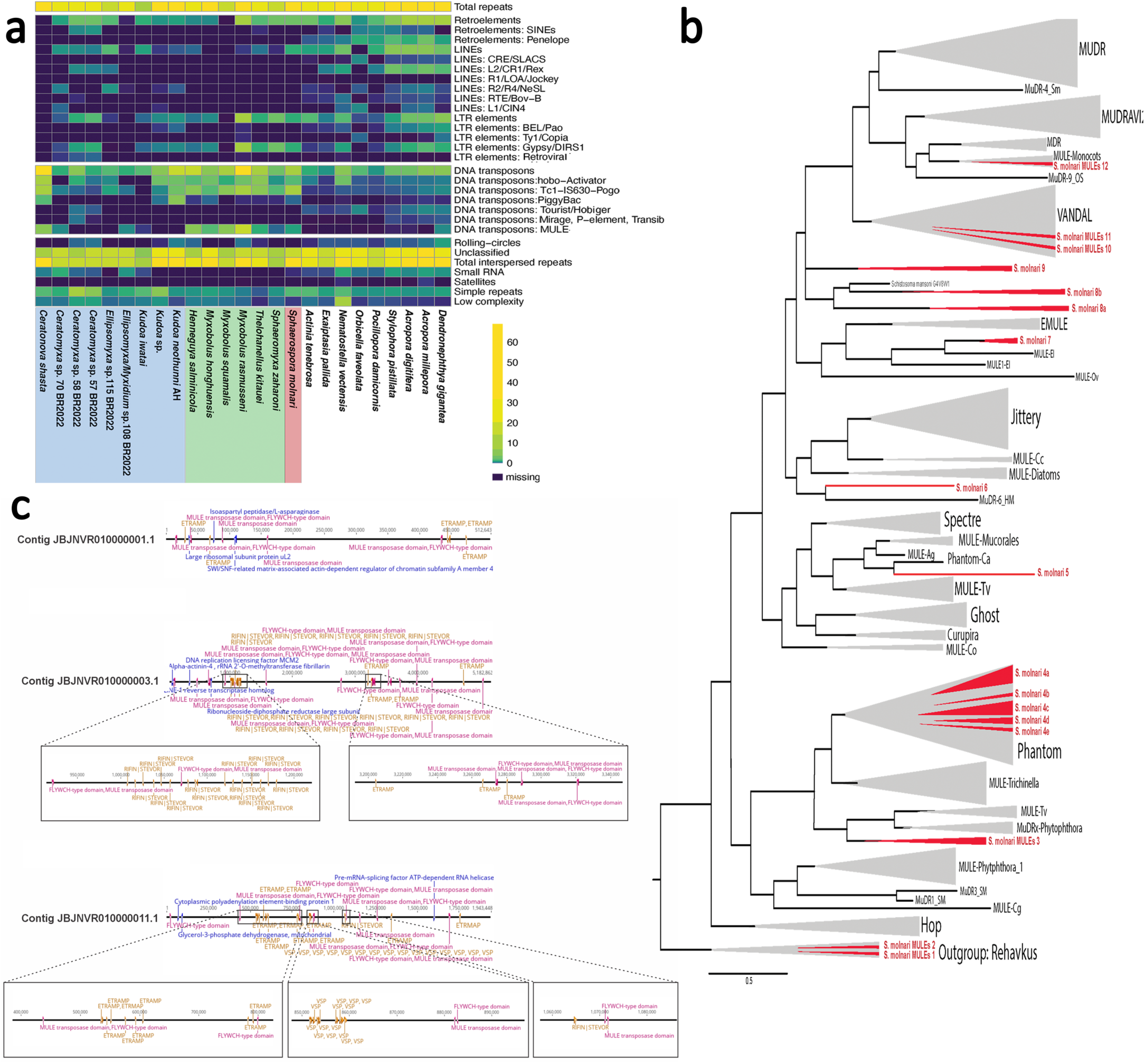
Repeat landscape and mutator-like elements (MULEs) in the genomes of myxozoans vs free-living cnidarians. **a.** Heatmap (RepeatMasker) illustrating important shifts in repeat family composition of myxozoan genomes, which show a pronounced reduction of retroelements (SINEs, LINEs, and most LTR elements), with *Gypsy/DIRS1* being the only retained group. While DNA transposons are also generally reduced, we observe a marked diversification of three categories: *PiggyBac*, *Tc1–IS630–Pogo*, and MULEs, predominantly in oligochaete-infecting myxozoans (green taxa). **b.** ML phylogenetic tree of MULE transposases of different eukaryotes and *S. molnari* (1,928 sequences) identified 12 independent origins of *S. molnari* MULEs (marked in red; 1-12), with several novel, phylogenetically basal lineages, and an important number of clades showing affinities to MULEs from different parasite groups (MuDRx – *Phytophthora*, MULE-Tv - *Trichomonas vaginalis,* MULE-Ei und EMULE – *Entamoeba* spp., Curupira – *Schistosoma mansoni*). The majority of *S. molnari* MULES belong to Phantom. Terminology according to Duperyron et al. [80]. Detailed tree in **Supplementary Figure 3. c.** Selected areas on three contigs of *S. molnari* which show clusters of MULES and ETRAMPs, RIFIN-STEVOR and VSPs indicating MULE-based propagation and diversification of these alien genes in *S. molnari*.

Comparative analyses revealed that Mutator-like elements (MULEs) are particularly diversified in oligochaete-infecting myxozoans, constituting up to 18.1% of all TEs, compared with only 0.6% on average in polychaete-infecting species and 0.06% on average in free-living cnidarians. In oligochaete-infecting myxozoans, MULE counts reached between 961 and 40,840 elements, occupying approx. 76x more genome space in these myxozoans than in the average free-living cnidarian.

MULEs are a significant superfamily of DNA transposons on account of their great transpositional activity and propensity for insertion in or near gene sequences, their consequent high mutagenic capacity, and their tendency to acquire host gene fragments [80]. MULEs encode a transposase that mediates their excision and integration, enabling the movement of DNA fragments within the genome [81]. Structurally, MULEs are characterized by terminal inverted repeats (TIRs) at both ends, which flank the transposase coding region. The transposase typically contains a conserved catalytic DDE motif and often includes an additional zinc-finger DNA-binding domain [80].

From an evolutionary perspective, the 141 MULE sequences identified in the genome of *S. molnari* can be ascribed to at least 12 independent phylogenetic origins (1-12) (**Figure 4b**), based on their transposase sequences. Several subclades represent novel, independent lineages that are unrelated to previously described eukaryotic MULEs (**Figure 4b**), many of them occupying basal positions within their respective subclades: Two sequences of *S. molnari* MULEs cluster with the most basal lineage, the Rehavkus group (1-2), previously described in annelids and insects. A cluster of 5 closely related *S. molnari* MULEs (3) branches sister to MULE-Tv from *Trichomonas vaginalis* and MULE-MuDRx from *Phytophthora*. This branch clusters basal to the large Phantom family to which most of *S. molnari*’s MULEs belong: 160 sequences are allocated to 5 clades (4a-e), with sister relationships to platyhelminths (a), molluscs (b), the parasitic crustacean fish parasite *Lepeophthirius salmonis* (c), parasitoid wasps (d), and an independent basal lineage (e) (**Figure 4b**). Phantom MULEs are structurally highly diverse and have been reported from a wide range of eukaryotic taxa, including many parasite lineages [82]. An additional *S. molnari* MULE clusters sister to Phantom-Ca from Ascomycota (5) (**Figure 4b**). The most basal lineage of the large clade containing the Jittery group (Rhodophyta) represents a cnidarian clade of MuDR-6 sequences from *Hydra magnipapillata* and an associated sequence from *S. molnari* (6). Basal to another major clade dominated by VANDAL and MuDR elements, four MULEs of *S. molnari* cluster with MULE-Ei and EMULEs of *Entamoeba* spp. (7). Two *S. molnari* branches of four and twelve MULE sequences (8a, 8b) cluster with a *Schistosoma mansoni* MULE (Curupira), one represents an independent lineage within the same subclade (9), and two fall within the VANDAL group (10-11), a non-TIR MULE family originally described from Brassicaceae plants [83]. Finally, two *S. molnari* MULEs cluster within the MULE-Monocots group (12) of Magnoliophyta. The diversity of phylogenetic lineages of MULEs recovered for *S. molnari* is surprising but may be explained by repeated capture, recombination, and reshuffling of endogenous and horizontally acquired genetic fragments.

The MULEs of maize are amongst the most mutagenic TEs, where large numbers of these elements lead to mutation frequencies 50 to 100 times that of their background [84]. This extremely high transposition frequency can exceed one new insertion per element per generation [85] and hence causes incomparable mutagenicity and recombination. Unlike typical MULEs that primarily move their own DNA, pack-MULEs, first described in maize, actively acquire fragments of cellular genes and rearrange them, contributing to gene evolution and genome plasticity. Not surprisingly MULEs have also been suggested to be involved in horizontal gene transfer (HGT) between organisms as distant as invertebrates and bats [86] or insects and mammals [87]. We screened *S. molnari* MULEs for embedded gene content and identified 99 genes in the pack-MULES of *S. molnari*. We detected numerous genes involved in central carbohydrate and energy metabolism, including glyceraldehyde-3-phosphate dehydrogenase, glycerol-3-phosphate dehydrogenase, enolase, glycogen phosphorylase, phosphofructokinase and fumarate hydratase. Together with genes encoding proteins required for mitochondrial maintenance (Lon protease, AFG3L2 (m-AAA protease), mitochondrial glycerol-3-phosphate dehydrogenase), these findings indicate that MULEs contribute to the amplification and diversification of genes involved in sugar-based energy metabolism and redox homeostasis in *S. molnari*. We further identified a substantial number of genes related to cytoskeleton, vesicle trafficking, transport proteins (dynein heavy chain, cynamin 2, alpha-actinin 4, gamma tubulin complex component 2, coatomer subunit epsilon) and membrane transporters (Na⁺/K⁺ ATPase α2, ABC transporter C family member 2, GLUT7 and monocarboxylate transporter 8). This suggests that *S. molnari* MULEs propagate genes related to intracellular transport, membrane trafficking and ion/metabolite uptake and discharge. In restricted areas of some genomic contigs, we found complexes of MULES with ETRAMPs, RIFIN-STEVOR and VSP as flanking genes, which appeared to be copied and pasted multiple times, in both directions in the genome (**Figure 4c**).

The extensive expansion of MULEs in myxozoans and the specific cargo of pack-MULEs suggests that these mutagenic elements significantly contribute to genomic change and adaptation, whether by self- or alien gene transposition, and we suggest they are likely the main drivers for the strongly accelerated evolutionary clock of myxozoans (Holzer et al. 2018). It would be of particular interest to investigate the frequencies of MULE insertion throughout the whole genome, their loci and flanking genes [88], to better understand the contribution of MULE transposons to the genetic variability, horizontal gene transfer and adaptive evolution of the Myxozoa.

### Gaining control over rapid genomic change: evolution of TFs from TEs

If the process of integration of TEs into the genome is followed by suppression of the self-propagation properties of these elements, we refer to molecular exaptation or domestication [89]. Domesticated, TE-derived gene regulators are an adaptive advantage [90] and provide a means of regulating the uncontrolled copy-paste activities of MULEs. The most important events of domestication are associated with the rise of biological novelty from the recruitment of TEs into transcription factors (TFs), as TFs are master regulators of gene expression in the Metazoa [91].

Similar to ctenophores, the earliest branching lineage of animals, which clusters basal to the Cnidaria [92], myxozoan parasites show an extraordinary expansion of the FLYWCH zinc finger DNA-binding site. In *M. rasmusseni*, FLYWCH is amongst the three most abundant protein domains in the genome [93]. In contrast, free-living cnidarians generally duplicated THAP, and hydrozoans a variety of different binding sites (ZBED, CENPB, HTH-Psq) [91]. Using DeepTFactor [94] and TransFacPred [95] we generated a catalog of MULE-derived TFs in the genome of *S. molnari* demonstrating the independent domestication of 13% of MULEs to TFs, via the FLYWCH DNA-binding site. The presence of transposon-like components within the protein-coding open reading frames (ORFs) of myxozoan TFs explains how the massive gene transposition in this group of parasites may be regulated while biological novelty is created.

In contrast to their free-living relatives, the TFs and transcriptional modulators of myxozoans but also those of some parasitic protists diversified considerably (e.g [96]). TFs may thus represent a major determinant shaping parasitism-related adaptation. Due to the general miniaturization of myxozoan genomes, only a small proportion of genes, mainly genes related to spore formation, maybe truly specific to a particular developmental stage of the parasite’s intrapiscine life cycle. Consequently, an extensive network of regulatory proteins needs to adjust gene expression on the transcriptional and post-transcriptional levels (review by e.g. [97]), which may explain the enlargement of the TF compartment in myxozoans.

## Conclusions

The Myxozoa is the oldest metazoan parasite group known to date. Myxozoans evolved alternation between invertebrate and vertebrate hosts and became obligate cnidarian parasites, they diversified successfully in their fish hosts, achieving a cosmopolitan distribution in most marine and freshwater habitats. Otherwise known as innocuous fish parasites, several species are recognized as pathogens of economically important wild and cultured fishes, with some taxa receiving special attention due to their emerging pathogen status in relation to climate change. Since the first myxozoan genome was published in 2014, myxozoan genomics has become a rapidly progressing field in this group. While several genomes have been published to date, the most basal lineage, Malacosporea, and the blood-proliferating lineage, *Sphaerospora sensu stricto*, had not been characterized so far. Considering that the myxozoan evolutionary rate is probably unmatched in speed amongst metazoans, it is essential to characterize the genomes of distantly related taxa and the variety of host exploitation and immune evasion strategies to identify common molecular tools for the design of multi-species approaches to treatments and vaccines. The phylogenetically limited number of published genome assemblies (16 species, only two out of four major branches), their lack of homology, and the high number of unknowns are considered the most substantial problems hampering progress in this field of research.

Using isolated blood parasites, we produced an Oxford Nanopore long read assembly of 40 near-chromosome-size contigs for *S. molnari,* the first myxozoan genome from the blood-proliferating sphaerosporid lineage. It is one of the most complete assemblies presently available for myxozoans, free from host contamination, with a high rate of gene discovery. The genomic features of *S. molnari* indicate that, despite clearly showing a reduced gene content due to parasitism, it represents a basal myxozoan genome, characterized by a single-circle mitochondrial genome, and by an unmatched completeness of eukaryotic core genes, metabolic pathways and functions, when compared with other species of myxozoans. We used this genomic database for comparative analyses of myxozoan gene groups that are either unique to myxozoans, or unique to the Sphaerospora lineage, have been duplicated and are hence likely to play a major role in myxozoan adaptation to a variety of parasitic lifestyles in fish. We discovered that DNA transposons belonging to the MULE superfamily are extremely common amongst the otherwise strongly reduced repetitive elements of myxozoans. Given the widespread occurrence of MULEs especially in oligochaete-infecting myxozoan genomes and the predominant presence of gene fragments of highly diversified gene families within pack-MULEs and in MULE-surrounding loci of *S. molnari* we hypothesize that gene fragment acquisition represents the most important mechanism for the evolution of new genes in this parasite group. MULEs may also be key to the transfer and successful establishment of taxonomically restricted alien genes from other parasites. We were able to prove the domestication of MULEs and their recruitment into transcription factors. Based on the massive duplication and diversification of these elements in myxozoan genomes, we hypothesize that MULEs and their derived TFs are the main drivers of genomic change and adaptation in the Myxozoa, and they most likely are the reason for their rapid evolutionary clock. Myxozoans further expanded genes coding for proteins that can modulate fish immune responses such as e.g. arginase repressors or functional domains for sequestering TGF-β in the extracellular matrix. As a blood parasite, *S. molnari* is particularly exposed to cellular and humoral immune responses, including specific antigenic recognition and immunoglobulin-based targeted elimination. As a unique immune evasion mechanism, the genome of *S. molnari* exhibits two types of variable surface proteins, with structural identity to VSPs of diplomonads and homology to VSAs of *Plasmodium*. It is likely that these allow *S. molnari* to vary its antigenic profile during its development in the fish, thereby remaining undiscovered by the hosts’ immune system for extended periods of time. Furthermore, highly charged surface proteins with homologous to ETRAMPs of *Plasmodium* possibly allow for efficient exploitation of target host cells (erythrocytes) by promoting attachment. It is likely that these genes were acquired by lateral gene transfer via MULEs, and they may well represent parasitic adaptations unique not only to this species but to the whole lineage of *Sphaerospora sensu stricto*, promoting survival and nutrition in the blood.

Our findings emphasize the importance of myxozoans for research as they demonstrate unique capabilities for adapting and acquiring new genes and genetic mechanisms, fueling the evolution of parasitism at the base of the Metazoa. The extensive high-quality genomic resource for *S. molnari* together with the findings from our targeted bioinformatic comparative exploitation build a solid base for future studies aiming at understanding the evolutionary origins and function of the highlighted parasite genes, as well as the remarkable mechanisms of gene acquisition and modification discovered in this study.

## Methods

### Sample collection, DNA isolation and sequencing

High molecular weight DNA for genome sequencing was isolated from *S. molnari* proliferative, presporogonic blood stages. *S. molnari* blood stages were collected from a laboratory line that has been transferred (4 + years) from fish to fish by intraperitoneal injection of parasites into specific parasite-free (SPF) common carp (*Cyprinus carpio*). To ensure minimum host contamination, parasites were isolated from infected fish blood via ion-exchange chromatography [19]. Isolated blood stages from 5 fish were processed with the Qiagen blood and tissue DNA extraction kit following manufacturer’s instructions. The Illumina sequencing library was created using the Nextera DNA Flex Library Prep Kit according to the manufacturers’ protocol (Illumina Inc. San Diego, CA, USA) and the quality was checked using the Agilent Bioanalyzer 2100 High Sensitivity DNA Kit (Agilent, Amstelveen, The Netherlands). The Illumina library was then sequenced using the NovaSeq6000 (2X150 nt) platform. Image analysis and basecalling were done by the Illumina pipeline. Nanopore reads were generated using the Oxford Nanopore Technologies platform (ONT, Oxford, UK). Library preparation was done with the SQK-LSK109 1D ligation kit (ONT, Oxford, UK). The ONT library was sequenced on a PromethION R9.4.1 flowcell (FLO-PRO002). Base-calling was done with GuPPy v1.6.0 [98], base-called reads were used for further processing and assembly.

### Genome assembly and functional annotation

Illumina reads were adapter- and quality-filtered using BBMap and seqtk/trimfq ([99]; https://github.com/lh3/seqtk). Nanopore reads were trimmed with Porechop v0.2.4 (https://github.com/rrwick/Porechop) and quality filtered with NanoFilt (https://doi.org/10.1093/bioinformatics/bty149) using q-value ≥ 7. To separate host from parasite sequences, nanopore reads were aligned to the *Cyprinus carpio* reference genome (PRJNA73579) with MiniMap2 [100], and the longest non-aligned reads were assembled *de novo* using multiple rounds of Flye v2.6 [101]. The final 40 Mb assembly, based on 35x coverage of the longest nanopore reads, was polished once with Racon [102], and subsequently with three Pilon v.1.23 [103] rounds, using the Illumina reads.

Gene prediction used an *ab initio* GeneMark-ET v4.61 [104] model to generate a preliminary gene set, which was then used to train AUGUSTUS v3.3.2 [105] gene models. Proteins from *Thelohanellus kitauei* (Genbank accession number GCA_000827895.1) were incorporated as hints, resulting in 14,957 predicted protein-coding genes. Structural genome characteristics were assessed using AGATe3 v1.13.0 (10.5281/zenodo.3552717). Functional annotation of predicted proteins followed a five-step pipeline: BLAST searches against (1) Uniprot/Swissprot, (2) EggNOG, (3) KEGG databases; (4) domain identification using HMMER3 and the Pfam-A database, (5) prediction of signal peptides and transmembrane domains with SignalP 5.0 ([106,107]) and TMHMM v 2.0 [108] tools. Annotation outputs from all databases were integrated into a .gff file. BUSCO v5.3.2 [109] and metazoa_odb10 was used for assessment of the gene content of the assembly.

### Chromosome visualization

Blood stages of *S. molnari* isolated by ion-exchange chromatography in RPMI medium (Gibco, Life Technologies), were enriched with 15% heat-inactivated fetal bovine serum (Biosera Europe, France) and 1 ml GlutaMAX™ (ThemoFisher, USA). To arrest nuclear divisions in metaphase, 0,01% colchicine was added to the culture prior to incubation at 21 °C overnight. The following day, blood stages received hypotonic treatment with 0,56 % KCl and were fixed in methanol:acetic acid (3:1), for 5 min. Parasites were spread on superfrost slides in 60 % acetic acid, dehydrated and dried. We used a universal fluorescence *in situ* hybridization (FISH) protocol [110] for mapping 18S rDNA and repetitive telomere sequences. The 18S probe was obtained by PCR from extracted parasite DNA using *S. molnari* specific primers [111]; SmSSU487F 5’-GCC TCT CCA CCT GTG TAT G-3’ and SmSSU1307R 5’-ACC GTG AGC CAC GCG TAA TG-3’) and biotin-16-dUTP (Jena Bioscience, Germany) while the biotin-labelled generic metazoan telomeric probe [TTAGGG]n [112] was obtained from Eurogentec (Seraing, Belgium). Briefly, slides were pre-treated with 100 μg/ml proteinase K (Serva, Heidelberg, Germany). Post hybridization and washing [110], probe hybridization to target DNA in chromosomes was visualized with Cy3-conjugated streptavidin (JacksonImmunoRes. Labs. Inc., Burlingame, CA, USA). Slides stained and mounted with an antifade medium containing DAPI. Chromosomes were observed and photographed on a confocal microscope (Olympus FV3000). TelFinder (https://github.com/bio-tombow/TelFinder) was used to identify telomeric regions bioinformatically.

### Phylogenomic analysis

The newly generated genome assembly of *S. molnari* and published genomic and transcriptomic datasets of ten myxozoans, fourteen other free-living cnidarians, and a sponge (outgroup) was used for phylogenomic analyses, in total comprising twenty-five species. Using BlastP (e-value set to e^-10^) this large dataset was screened for 78 ribosomal protein-coding genes from a publicly available dataset (http://www.treebase.org, accession number S10436; [113]). Candidate sequences for all taxa were obtained for 77 of the 78 genes. For each gene, the top five sequences were retrieved from the BlastP output. Selected sequences were then filtered using PREQUAL v1.02 [114] with default options to remove non-homologous regions present in low-quality sequences. Sequences were aligned with MAFFT LINSI v7.453 [115], using default options. Aligned sequences were subjected to partial filtering using Divvier (https://github.com/simonwhelan/Divvier) with divvygap and mincol set to 4. Ambiguously aligned regions were excluded with trimAl v 1.4 [116], using a gap threshold of 1%. Initial trees were computed with IQ-TREE v. 1.6.12, using C60+LG+G+f model, and 100 PMSF bootstrap replicates [117]. All alignments were manually inspected, questionable sequences were removed. The remaining orthologs were realigned following the previously described steps and concatenated into a supermatrix using catfasta2phyml (https://github.com/nylander/catfasta2phyml). The resulting alignment comprised 12,809 amino acid positions. The final tree was calculated using IQ-TREE v. 1.6.12 with previously described parameters.

### Mitochondrial genome analyses

Adapter- and quality-filtered Illumina reads were *de novo* assembled with IDBA-UD v 1.1.3 [118]. The *cox1* amino-acid sequence of *Kudoa iwatai* (GenBank accession number BAR94716.1) was used as tblastn query, and resulting hits served as seed sequences for NOVOplasty v. 4.1 [119], in “mito” mode (read length of 150 bp and genome range 12-40 kb), producing a single 15,106 bp contig. Read mapping confirmed circularity and coverage. Using this contig as query, Homologous long reads were extracted by blastn, yielding 6,676 mitochondrial reads (41 Mb; N50 8,047 bp), including 116 reads >15,106 bp, of which 25 were 13.8 kb concatemers. BLAST analyses revealed a 952 bp short-read miss-assembly in the 15,106 bp contig. The 13.8 kb long read was corrected with RACON v1.3.3 [102] and polished twice with Pilon, generating a 14,015 bp consensus mt genome (GenBank accession number PQ893892).

Mt genome annotation followed multiple approaches. ORFs were identified with the Geneious Prime v2023.2.1 “Find ORFs” tool [45] and annotated with MITOS2 [120] using translation **Table 4**. ORFs were aligned to orthologous myxozoan sequences, and gene boundaries were defined by minimizing overlap and non-coding sequence while maximizing similarity. Transmembrane domains were predicted using Geneious Prime “Predict Transmembrane Regions v1.0.2.” ORFs on the reverse strand or nested within other genes were excluded. Only three protein-coding genes were recovered; additional searches with MFannot [121] and HMMER3 v3.3.2 (http://hmmer.org) identified no further genes. Ribosomal genes (*rnl*, *rns*) were annotated with MITOS2 and validated by RNA-seq read mapping (SRX23359792–94) following Takeuchi et al. [48]. These mappings also confirmed the identified ORFs. tRNA searches using tRNAscan-SE [122], RNAweasel [121], RFAM [123], and MITOS2 (accessed Nov 2024) did not yield manually verifiable tRNAs. Repeats were identified using Repeat Finder v1.0.1 in Geneious Prime as described in Sandberg et al. [45].

### Pathway analyses

The completeness of metabolic pathways in the *S. molnari* genome assembly was assessed using the Kyoto Encyclopedia of Genes and Genomes (KEGG) framework. Predicted gene models derived from the *S. molnari* genome were functionally annotated using the KEGG Automatic Annotation Server (KAAS; https://www.genome.jp/tools/kaas/). Orthologous relationships were assigned by mapping protein sequences to KEGG Orthology (KO) identifiers using the single-directional best-hit method against a representative eukaryotic dataset. The resulting KO annotations were used to reconstruct metabolic and cellular pathways by mapping annotated genes onto KEGG reference pathway maps. Comparative approaches included KEGG analyses of the genomes of seven other myxozoan species (**Supplementary figure 2**). Pathway completeness was visualized using the KEGG Mapper tool (www.genome.jp/kegg/tool/map_pathway1.html).

### Gene orthology, lineage-specific genes and duplication events

We used OrthoFinder v2.3.10 [124] to infer orthologous groups (orthogroups) among gene models from 11 free-living cnidarians and 9 myxozoan species, using *Amphimedon queenslandica* as the outgroup. Orthogroups were functionally annotated by BLAST searches of representative sequences against the UniProt/Swiss-Prot database. *S. molnari*–specific and myxozoan-specific orthogroups were identified from the “Orthogroups.GeneCount” table produced by OrthoFinder. Orthogroups were selected using a presence/absence criterion: (i) orthogroups containing zero genes in free-living cnidarians but ≥1 gene in ≥2 myxozoans were classified as myxozoan-specific; (ii) orthogroups containing zero genes in myxozoans but ≥1 gene in *S. molnari* were classified as *S. molnari*–specific. For the myxozoan-specific subset, we further prioritized orthogroups represented in at least four of the nine myxozoan species, ensuring robust lineage-specific signal.

To investigate potential sources of functional innovation, we examined genes and domains that had undergone extensive duplication (≥5 copies). Gene copy numbers were derived from the annotated *S. molnari* gene models by clustering predicted proteins using sequence similarity–based approaches (BLASTP followed by Markov clustering) to identify paralogous groups. Gene families or domains exceeding the ≥5-copy threshold were extracted for downstream analysis with custom scripts. For each duplicated family, we inspected gene architecture, domain composition, and genomic context to evaluate whether duplication events might contribute to neofunctionalization or subfunctionalization. This analysis allowed us to highlight candidate gene families potentially involved in lineage-specific adaptations.

### Identification of VSPs

Putative VSPs were identified following the following pipeline [65]: (1) identification of cysteine rich proteins (at least 5% of cysteine in a transcript), (2) identification of a transmembrane (TM) domain using TMHMM v2.0, (3) identification of at least 6 motifs of CXC or CXXC, (4) identification of conserved motifs at the C terminus, (5) observation of sequence length between TM domain and C terminus.

### Identification of repetitive elements and characterization of MULEs

Repetitive elements in the genome assembly of *S. molnari* were identified by: (1) building a repeat library using RepeatModeler2 v. 1.0.7 [125], and (2) running RepeatMasker version 4.0.7 [126] by using the custom library produced by RepeatModeler2 in the previous step. The resulting sets of repetitive elements were annotated by ABBlast/WUBlast searches against Master RepeatMasker database (dc20170127-rb20170127) bundled with RepeatMasker. A similar pipeline was used to identify the repeat elements in other publicly available cnidarian genomes, and results were plotted as a heatmap for comparison.

Phylogenetic relationships between the MULE transposases found in the genome of *S. molnari* and those of other eukaryotes were inferred using a maximum-likelihood (ML) approach. Using MAFFT [115] and the FFT-NS-i strategy with --maxiterate 1000 and the BLOSUM62 scoring matrix (implemented in Geneious Prime v2023.2.1), 1,950 amino-acid sequences were aligned. ML phylogenies were reconstructed in RAxML [127] using the PROTGAMMAWAG model (WAG substitution matrix with Γ-distributed rate heterogeneity; alpha parameter estimated). *S. molnari* MULEs were validated, screened and annotated using Censor [128]. Pack-MULEs were detected by screening contigs for terminal inverted repeats (TIRs) with EMBOSS einverted [129]. BLAST hits fully contained within TIR spans were retained, and corresponding gene regions were extracted using SAMtools [130]. Candidate elements were screened for protein-coding content using DIAMOND BLASTX [131] against Swiss-Prot. Regions with host-gene matches underwent refined TIR detection to define element boundaries, with emphasis on elements carrying metabolic genes selected for detailed analysis.

## Data availability

All *S. molnari* Oxford Nanopore and Illumina raw sequence data have been deposited in the National Center for Biotechnology Information (NCBI) (SRA database under BioProject accession PRJNA1193866).

## Supplementary Figures

**Supplementary Figure 1.** RNA reads mapped to the mitochondrial genome of *S. molnari*, confirming all gene loci, but showing insufficient read-length coverage over the *rns* region resulting in a tentative annotation of *rns* based on MITOS2 output.

**Supplementary Figure 2.** Metabolic heatmap (KEGG Mapper) comparing KEGG pathway completeness across nine myxozoan species. *S. molnari* stands out as the most metabolically complete and least reduced myxozoan included in the analysis.

**Supplementary Figure 3.** ML phylogenetic tree (MAFFT implemented in Geneious Pro) of eukaryotic MULE transposases showing the phylogenetic origins (12) of *S. molnari* MULEs and their relation with those of other taxa. Terminology according to [80].

## Acknowledgements

We highly appreciate training and advice in chromosome FISH methods by Prof. František Marec and Dr. Magda Zrzavá (Department of Molecular Biology and Genetics (University of South Bohemia)/Institute of Entomolgy (Biology Centre of the Czech Academy of Sciences).

## Author Information

Authors and Affiliations

## Contributions

AK and ASH designed the research. ASH generated the parasite material. RPD supervised the whole-genome assembly work at Future Genomics Technologies. AK, MW, RPD, IF, ERF, GAB and AS analyzed the data. ASH wrote the article with input from the other authors. All authors reviewed the manuscript.

## Corresponding authors

Correspondence to Astrid S. Holzer.

## Ethics declaration

### Ethics approval

Experimental infection and animal handling in this study was carried out in accordance with the Animal Protection Law of the Czech Republic No. 246/1992 Coll., ethics approval No. 88/2019, and protocols approved by the responsible committee of the Institute of Parasitology, Biology Centre of the Czech Academy of Sciences (PAU, BC CAS), compliant with with the relevant European guidelines on animal welfare (Directive 2010/63/EU on the protection of animals used for scientific purposes) and the recommendations of the Federation of Laboratory Animal Science Associations.

### Consent for publication

Not applicable.

## References

[1] Ahuja N, Cao X, Schultz DT, et al. Giants among Cnidaria: Large nuclear genomes and tearranged mitochondrial genomes in siphonophores. Genome Biol Evol. 2024;16(3).

[2] Alama-Bermejo G, Holzer AS. Advances and discoveries in myxozoan genomics. Trends Parasitol. 2021;37(6):552–568.

[3] Hartikainan H, Doonan L, Hartigan A, et al. Comparative genomics of species in an early diverging myxozoan clade - the Malacosporea. Abstract book of the 20th International Conference of Diseases of Fish and Shellfish. 2021;102.

[4] Guo Q, Atkinson SD, Xiao B, et al. A myxozoan genome reveals mosaic evolution in a parasitic cnidarian. BMC Biol. 2022;20(1):51.

[5] Muthye VR, Leon Coria A, Liu H, et al. The highly repetitive genome of *Myxobolus rasmusseni*, an emerging myxozoan parasite of fathead minnows. Genome Biol Evol. 2024;16(11).

[6] Slyusarev GS, Starunov V V., Bondarenko AS, et al. Extreme genome and nervous system streamlining in the invertebrate parasite *Intoshia variabili*. Current Biology. 2020;30(7):1292–1298.e3.

[7] Poulin R, Randhawa HS. Evolution of parasitism along convergent lines: from ecology to genomics. Parasitology. 2015;142(S1):S6–S15.

[8] Chang ES, Neuhof M, Rubinstein ND, et al. Genomic insights into the evolutionary origin of Myxozoa within Cnidaria. Proceedings of the National Academy of Sciences. 2015;112(48):14912–14917.

[9] Neverov AM, Panchin AY, Mikhailov K V., et al. Apoptotic gene loss in Cnidaria is associated with transition to parasitism. Sci Rep. 2023;13(1):8015.

[10] Graham AM, Barreto FS. Myxozoans (Cnidaria) do not retain key oxygen-sensing and homeostasis toolkit genes. Genome Biol Evol. 2023;15(1).

[11] Yahalomi D, Atkinson SD, Neuhof M, et al. A cnidarian parasite of salmon (Myxozoa: *Henneguya*) lacks a mitochondrial genome. Proceedings of the National Academy of Sciences. 2020;117(10):5358–5363.

[12] Holland JW, Holzer AS. Myxozoan Research Forum 2021 - the ‘MyxoMixer’: Advances, methods, and problems yet to be solved in myxozoan research. Bull Eur Assoc Fish Pathol. 2022;41(5).

[13] Bartošová P, Fiala I, Jirků M, et al. *Sphaerospora sensu stricto*: Taxonomy, diversity and evolution of a unique lineage of myxosporeans (Myxozoa). Mol Phylogenet Evol. 2013;68(1):93–105.

[14] Hartigan A, Kosakyan A, Pecková H, et al. Transcriptome of *Sphaerospora molnari* (Cnidaria, Myxosporea) blood stages provides proteolytic arsenal as potential therapeutic targets against sphaerosporosis in common carp. BMC Genomics. 2020;21(1):404.

[15] Korytář T, Chan JTH, Vancová M, et al. Blood feast: Exploring the erythrocyte-feeding behaviour of the myxozoan *Sphaerospora molnari*. Parasite Immunol. 2020;42(8).

[16] Anderson HL, Brodsky IE, Mangalmurti NS. The evolving erythrocyte: Red blood cells as modulators of innate immunity. The Journal of Immunology. 2018;201(5):1343–1351.

[17] Majstorović J, Kyslík J, Klak K, et al. Erythrocytes of the common carp are immune sentinels that sense pathogen molecular patterns, engulf particles and secrete pro-inflammatory cytokines against bacterial infection. Front Immunol. 2024;15.

[18] Korytář T, Wiegertjes GF, Zusková E, et al. The kinetics of cellular and humoral immune responses of common carp to presporogonic development of the myxozoan *Sphaerospora molnari*. Parasit Vectors. 2019;12(1):208.

[19] Born-Torrijos A, Kosakyan A, Patra S, et al. Method for isolation of myxozoan proliferative stages from fish at high yield and purity: An essential prerequisite for *in vitro*, *in vivo* and genomics-based research developments. Cells. 2022;11(3):377.

[20] Holzer AS, Bartošová-Sojková P, Born-Torrijos A, et al. The joint evolution of the Myxozoa and their alternate hosts: A cnidarian recipe for success and vast biodiversity. Mol Ecol. 2018;27(7):1651–1666.

[21] Yuan J, Li J, Yong J, et al. A telomere-to-telomere genome assembly of koi carp (*Cyprinus carpio*) using long reads and Hi-C technology. Gigascience. 2025;14.

[22] Waterhouse RM, Tegenfeldt F, Li J, et al. OrthoDB: A hierarchical catalog of animal, fungal and bacterial orthologs. Nucleic Acids Res. 2013;41(D1):D358–D365.

[23] Starcevic A, Figueredo RTA, Naldoni J, et al. Long-read metagenomic sequencing negates inferred loss of cytosine methylation in Myxosporea (Cnidaria: Myxozoa). Gigascience. 2025;14.

[24] Weber CC, Paulini M, Blaxter ML. Myxozoan parasite genomes assembled from contaminated host data reveal extensive gene order conservation and rapid sequence evolution. G3: Genes, Genomes, Genetics. 2025;15(7).

[25] Vacarizas J, Taguchi T, Mezaki T, et al. Cytogenetic markers using single-sequence probes reveal chromosomal locations of tandemly repetitive genes in scleractinian coral *Acropora pruinosa*. Sci Rep. 2021;11(1):11326.

[26] Kawakami R, Taguchi T, Vacarizas J, et al. Karyotypic analysis and isolation of four DNA markers of the scleractinian coral *Favites pentagona* (Esper, 1795) (Scleractinia, Anthozoa, Cnidaria). Comp Cytogenet. 2022;16(1):77–92.

[27] Georgevitch J. Über Diplo- und Haplophase im Entwicklungskreise der Myxosporidien. Arch Protistenkunde. 1935;84:419–428.

[28] Georgevitch J. Über Diplo- und Haplophase im Entwickungskreise des Myxosporids *Zschokkella rovignensis* Nemeczek. Arch Protistenkunde. 1936;87:151–154.

[29] Noble ER. Nuclear cycles in the life history of the protozoan genus *Ceratomyxa*. J Morphol. 1941;69(3):455–479.

[30] Noble ER. Nuclear cycles in the protozoan parasite *Myxidium gasterostei* n. sp. J Morphol. 1943;73(2):281–295.

[31] Atkinson SD, O’Neil ST, Meyer E, et al. First look at the genome of *Ceratomyxa shasta*, a myxozoan parasite of salmonids). Abstract book of the 19th International Conference of Diseases of Fish and Shellfish. 2013;68.

[32] Davids HS. The structure and development of a myxosporidian parasite of the sequeteague *Cynoscion regalis*. J Morphol. 1916;27:333–377.

[33] Behlar K. Die Formwechsel der Protistenkerne. Eine vergleichende morphologische Studie. Ergebnisse und Fortschritte der Zoologie. 1926;6:235–254.

[34] Weiser J. Parasites of freshwater fish. Věstník Československé Zoologické Společnosti. 1949;13:364–371.

[35] Hallett S, Bartholomew J. Application of a real-time PCR assay to detect and quantify the myxozoan parasite *Ceratomyxa shasta* in river water samples. Dis Aquat Organ. 2006;71:109–118.

[36] Alama-Bermejo G, Šíma R, Raga JA, et al. Understanding myxozoan infection dynamics in the sea: Seasonality and transmission of *Ceratomyxa puntazzi*. Int J Parasitol. 2013;43(9):771–780.

[37] Traut W, Szczepanowski M, Vítková M, et al. The telomere repeat motif of basal Metazoa. Chromosome Research. 2007; doi: 10.1007/s10577-007-1132-3.

[38] Scherf A. *Plasmodium* telomeres: a pathogen’s perspective. Curr Opin Microbiol. 2001;4(4):409–414.

[39] Fiala I. The phylogeny of Myxosporea (Myxozoa) based on small subunit ribosomal RNA gene analysis. Int J Parasitol. 2006;36(14):1521–1534.

[40] Fiala I, Bartošová-Sojková P, Whipps CM. Classification and Phylogenetics of Myxozoa. Myxozoan Evolution, Ecology and Development. Cham: Springer International Publishing; 2015. p. 85–110.

[41] Molnár K, El-Mansy A, Székely C, et al. Experimental identification of the actinosporean stage of *Sphaerospora renicola* Dykova & Lom 1982 (Myxosporea: Sphaerosporidae) in oligochaete alternate hosts. J Fish Dis. 1999;22(2):143–153.

[42] Novosolov M, Yahalomi D, Chang ES, et al. The phylogenetic position of the enigmatic, *Polypodium hydriforme* (Cnidaria, Polypodiozoa): Insights from mitochondrial genomes. Genome Biol Evol. 2022;14(8).

[43] Chang ES, Neuhof M, Rubinstein ND, et al. Genomic insights into the evolutionary origin of Myxozoa within Cnidaria. Proceedings of the National Academy of Sciences. 2015;112(48):14912–14917.

[44] Patra S, Bartošová-Sojková P, Pecková H, et al. Biodiversity and host-parasite cophylogeny of *Sphaerospora* (*sensu stricto*) (Cnidaria: Myxozoa). Parasit Vectors. 2018;11(1):347.

[45] Sandberg TOM, Yahalomi D, Bracha N, et al. Evolution of myxozoan mitochondrial genomes: insights from myxobolids. BMC Genomics. 2024;25(1):388.

[46] Yahalomi D, Haddas-Sasson M, Rubinstein ND, et al. The multipartite mitochondrial genome of *Enteromyxum leei* (Myxozoa): Eight gast-rvolving megacircles. Mol Biol Evol. 2017;34(7):1551–1556.

[47] Feng H, Lv S, Li R, et al. Mitochondrial genome comparison reveals the evolution of cnidarians. Ecol Evol. 2023;13(6).

[48] Takeuchi F, Sekizuka T, Ogasawara Y, et al. The mitochondrial genomes of a myxozoan genus *Kudoa* are extremely divergent in Metazoa. PLoS One. 2015;10(7):e0132030.

[49] Ling F, Niu R, Hatakeyama H, et al. Reactive oxygen species stimulate mitochondrial allele segregation toward homoplasmy in human cells. Mol Biol Cell. 2016;27(10):1684–1693.

[50] Tan J, Dong X, Liu H. Mitochondrial DNA is a sensitive surrogate and oxidative stress target in oral cancer cells. PLoS One. 2024;19(9):e0304939.

[51] Meena M, Divyanshu K, Kumar S, et al. Regulation of L-proline biosynthesis, signal transduction, transport, accumulation and its vital role in plants during variable environmental conditions. Heliyon. 2019;5(12):e02952.

[52] Rath M, MÃ¼ller I, Kropf P, et al. Metabolism via arginase or nitric oxide synthase: Two competing arginine pathways in macrophages. Front Immunol. 2014;5.

[53] Wentzel AS, Janssen JJE, de Boer VCJ, et al. Fish macrophages show distinct metabolic signatures upon polarization. Front Immunol. 2020;11.

[54] Holzmuller P, Geiger A, Nzoumbou-Boko R, et al. Trypanosomatid infections: How do parasites and their excreted–secreted factors modulate the inducible metabolism of L-arginine in macrophages? Front Immunol. 2018;9.

[55] Schofield PJ, Edwards MR, Matthews J, et al. The pathway of arginine catabolism in *Giardia intestinalis*. Mol Biochem Parasitol. 1992;51(1):29–36.

[56] Stadelmann B, Merino MC, Persson L, et al. Arginine consumption by the intestinal parasite *Giardia intestinalis* reduces proliferation of intestinal epithelial cells. PLoS One. 2012;7(9):e45325.

[57] Bailey C, Strepparava N, Ros A, et al. It’s a hard knock life for some: Heterogeneity in infection life history of salmonids influences parasite disease outcomes. Journal of Animal Ecology. 2021;90(11):2573–2593.

[58] Gobert AP, Daulouede S, Lepoivre M, et al. L-arginine availability modulates local nitric oxide production and parasite killing in experimental trypanosomiasis. Infect Infectmmun. 2000;68(8):4653–4657.

[59] Gutiérrez-Kobeh L, González JR, Vázquez-López R, et al. Signaling Pathways Targeted by Protozoan Parasites to Inhibit Apoptosis. Current Understanding of Apoptosis - Programmed Cell Death. InTech; 2018.

[60] Arabiotorre A, Bankaitis VA, Grabon A. Regulation of phosphoinositide metabolism in Apicomplexan parasites. Front Cell Dev Biol. 2023;11.

[61] Freitas TC, Jung E, Pearce EJ. TGF-β Signaling Controls Embryo Development in the Parasitic Flatworm Schistosoma mansoni. PLoS Pathog. 2007;3(4):e52.

[62] Mukundan A, Byeon C-H, Hinck CS, et al. Convergent evolution of a parasite-encoded complement control protein-scaffold to mimic binding of mammalian TGF-β to its receptors, TβRI and TβRII. Journal of Biological Chemistry. 2022;298(6):101994.

[63] Robertson IB, Rifkin DB. Regulation of the Bioavailability of TGF-β and TGF-β-Related Proteins. Cold Spring Harb Perspect Biol. 2016;8(6):a021907.

[64] Mowatt MR, Aggarwal A, Nash TE. Carboxy-terminal sequence conservation among variant-specific surface proteins of Giardia lamblia. Mol Biochem Parasitol. 1991;49(2):215–227.

[65] Wiśniewska MM, Salomaki ED, Silberman JD, et al. Expanded gene and taxon sampling of diplomonads shows multiple switches to parasitic and free-living lifestyle. BMC Biol. 2024;22(1):217.

[66] Adam RD, Nigam A, Seshadri V, et al. The Giardia lamblia vsp gene repertoire: characteristics, genomic organization, and evolution. BMC Genomics. 2010;11(1):424.

[67] Braden LM, Rasmussen KJ, Purcell SL, et al. Acquired Protective Immunity in Atlantic Salmon Salmo salar against the Myxozoan Kudoa thyrsites Involves Induction of MHIIβ ^+^ CD83 ^+^ Antigen-Presenting Cells. Infect Immun. 2018;86(1).

[68] Morris D, Morris DC, Adams A. Development and release of a malacosporean (Myxozoa) from Plumatella repens (Bryozoa: Phylactolaemata). J Aquat Anim Health. 1997;9:265–272.

[69] Cheng Q, Cloonan N, Fischer K, et al. stevor and rif are *Plasmodium falciparum* multicopy gene families which potentially encode variant antigens. Mol Biochem Parasitol. 1998;97(1–2):161–176.

[70] Niang M, Bei AK, Madnani KG, et al. STEVOR Is a *Plasmodium falciparum* erythrocyte binding protein that mediates merozoite invasion and rosetting. Cell Host Microbe. 2014;16(1):81–93.

[71] Saito F, Hirayasu K, Satoh T, et al. Immune evasion of *Plasmodium falciparum* by RIFIN via inhibitory receptors. Nature. 2017;552(7683):101–105.

[72] Sakoguchi A, Chamberlain SG, Mørch AM, et al. RIFINs displayed on malaria-infected erythrocytes bind KIR2DL1 and KIR2DS1. Nature. 2025;643(8074):1363–1371.

[73] Chew M, Ye W, Omelianczyk RI, et al. Selective expression of variant surface antigens enables *Plasmodium falciparum* to evade immune clearance in vivo. Nat Commun. 2022;13(1):4067.

[74] Spielmann T, Fergusen DJP, Beck H-P. ETAMPs, a New *Plasmodium falciparum* gene family coding for developmentally regulated and highly charged membrane proteins located at the parasite–host vell interface. Mol Biol Cell. 2003;14(4):1529–1544.

[75] Korytář T, Chan JTH, Vancová M, et al. Blood feast: Exploring the erythrocyte-feeding behaviour of the myxozoan *Sphaerospora molnari*. Parasite Immunol. 2020;42(8).

[76] Nakamura K, Oshima T, Morimoto T, et al. Sequence-specific error profile of Illumina sequencers. Nucleic Acids Res. 2011;39(13):e90–e90.

[77] Thomas MC, Macias F, Alonso C, et al. The biology and evolution of transposable elements in parasites. Trends Parasitol. 2010;26(7):350–362.

[78] Lisch D. *Mutator* and *MULE* Transposons. Microbiol Spectr. 2015;3(2).

[79] Liu K, Wessler SR. Transposition of Mutator–like transposable elements (MULEs) resembles hAT and Transib elements and V(D)J recombination. Nucleic Acids Res. 2017;45(11):6644–6655.

[80] Dupeyron M, Singh KS, Bass C, et al. Evolution of Mutator transposable elements across eukaryotic diversity. Mob DNA. 2019;10(1):12.

[81] Feschotte C, Pritham EJ. DNA transposons and the evolution of eukaryotic genomes. Annu Rev Genet. 2007;41(1):331–368.

[82] Marquez CP, Pritham EJ. *Phantom*, a New Subclass of *Mutator* DNA Transposons Found in Insect Viruses and Widely Distributed in Animals. Genetics. 2010;185(4):1507–1517.

[83] Kapitonov V V., Jurka J. Molecular paleontology of transposable elements from Arabidopsis thaliana. Genetica. 1999;107(1–3):27–37.

[84] Walbot V. The Mutator transposable element family of Maize. Genet Eng (N Y). Boston, MA: Springer US; 1991. p. 1–37.

[85] Alleman M, Freeling M. The *Mu* transposable elements of maize: Evidence for transposition and copy number regulation during development. Genetics. 1986;112(1):107–119.

[86] Tang Z, Zhang H-H, Huang K, et al. Repeated horizontal transfers of four DNA transposons in invertebrates and bats. Mob DNA. 2015;6(1):3.

[87] Oliveira SG, Bao W, Martins C, et al. Horizontal transfers of Mariner transposons between mammals and insects. Mob DNA. 2012;3(1):14.

[88] Bradic M, Warring SD, Low V, et al. The Tc1/mariner transposable element family shapes genetic variation and gene expression in the protist *Trichomonas vaginalis*. Mob DNA. 2014;5(1):12.

[89] Gould SJ, Vrba ES. Exaptation—a missing term in the science of rorm. Paleobiology. 1982;8(1):4–15.

[90] Miller WJ, McDonald JF, Nouaud D, et al. Molecular domestication – more than a sporadic episode in evolution. Genetica. 1999;107(1–3):197–207.

[91] Mukherjee K, Moroz LL. Transposon-derived transcription factors across metazoans. Front Cell Dev Biol. 2023;11.

[92] Whelan N V., Kocot KM, Moroz TP, et al. Ctenophore relationships and their placement as the sister group to all other animals. Nat Ecol Evol. 2017;1(11):1737–1746.

[93] Muthye VR, Leon Coria A, Liu H, et al. The highly repetitive genome of Myxobolus sp., a myxozoan parasite of fathead minnows. 2024.

[94] Kim GB, Gao Y, Palsson BO, et al. DeepTFactor: A deep learning-based tool for the prediction of transcription factors. Proceedings of the National Academy of Sciences. 2021;118(2).

[95] Patiyal S, Tiwari P, Ghai M, et al. A hybrid approach for predicting transcription factors. Frontiers in Bioinformatics. 2024;4.

[96] Iyer LM, Anantharaman V, Wolf MY, et al. Comparative genomics of transcription factors and chromatin proteins in parasitic protists and other eukaryotes. Int J Parasitol. 2008;38(1):1–31.

[97] Bennink S, Pradel G. The molecular machinery of translational control in malaria parasites. Mol Microbiol. 2019;112(6):1658–1673.

[98] Sherathiya VN, Schaid MD, Seiler JL, et al. GuPPy, a Python toolbox for the analysis of fiber photometry data. Sci Rep. 2021;11(1):24212.

[99] Bushnell B, Rood J, Singer E. BBMerge – Accurate paired shotgun read merging via overlap. PLoS One. 2017;12(10):e0185056.

[100] Li H. Minimap2: pairwise alignment for nucleotide sequences. Bioinformatics. 2018;34(18):3094–3100.

[101] Kolmogorov M, Yuan J, Lin Y, et al. Assembly of long, error-prone reads using repeat graphs. Nat Biotechnol. 2019;37(5):540–546.

[102] Vaser R, Sović I, Nagarajan N, et al. Fast and accurate de novo genome assembly from long uncorrected reads. Genome Res. 2017;27(5):737–746.

[103] Walker BJ, Abeel T, Shea T, et al. Pilon: An integrated tool for comprehensive microbial variant detection and genome assembly improvement. PLoS One. 2014;9(11):e112963.

[104] Brůna T, Lomsadze A, Borodovsky M. GeneMark-EP+: eukaryotic gene prediction with self-training in the space of genes and proteins. NAR Genom Bioinform. 2020;2(2).

[105] Stanke M, Keller O, Gunduz I, et al. AUGUSTUS: ab initio prediction of alternative transcripts. Nucleic Acids Res. 2006;34(Web Server):W435–W439.

[106] Chen Y, Wan Y, Pei X, et al. GATA3 differentially regulates the transcriptome via zinc finger 2-modulated phase separation. Cell Rep. 2025;44(5):115702.

[107] Almagro Armenteros JJ, Tsirigos KD, Sønderby CK, et al. SignalP 5.0 improves signal peptide predictions using deep neural networks. Nat Biotechnol. 2019;37(4):420–423.

[108] Krogh A, Larsson B, von Heijne G, et al. Predicting transmembrane protein topology with a hidden markov model: application to complete genomes11Edited by F. Cohen. J Mol Biol. 2001;305(3):567–580.

[109] Manni M, Berkeley MR, Seppey M, et al. BUSCO: Assessing Genomic Data Quality and Beyond. Curr Protoc. 2021;1(12).

[110] Cabral-de-Mello DC, Marec F. Universal fluorescence in situ hybridization (FISH) protocol for mapping repetitive DNAs in insects and other arthropods. Molecular Genetics and Genomics. 2021;296(3):513–526.

[111] Eszterbauer E, Sipos D, Forró B, et al. Molecular characterization of Sphaerospora molnari (Myxozoa), the agent of gill sphaerosporosis in common carp Cyprinus carpio carpio. Dis Aquat Organ. 2013;104(1):59–67.

[112] Moyzis RK, Buckingham JM, Cram LS, et al. A highly conserved repetitive DNA sequence, (TTAGGG)n, present at the telomeres of human chromosomes. Proceedings of the National Academy of Sciences. 1988;85(18):6622–6626.

[113] Nesnidal MP, Helmkampf M, Bruchhaus I, et al. Compositional Heterogeneity and Phylogenomic Inference of Metazoan Relationships. Mol Biol Evol. 2010;27(9):2095–2104.

[114] Whelan S, Irisarri I, Burki F. PREQUAL: detecting non-homologous characters in sets of unaligned homologous sequences. Bioinformatics. 2018;34(22):3929–3930.

[115] Katoh K, Standley DM. MAFFT Multiple Sequence Alignment Software Version 7: Improvements in Performance and Usability. Mol Biol Evol. 2013;30(4):772–780.

[116] Capella-Gutiérrez S, Silla-Martínez JM, Gabaldón T. trimAl: a tool for automated alignment trimming in large-scale phylogenetic analyses. Bioinformatics. 2009;25(15):1972–1973.

[117] Nguyen L-T, Schmidt HA, von Haeseler A, et al. IQ-TREE: A fast and effective stochastic algorithm for estimating maximum-likelihood phylogenies. Mol Biol Evol. 2015;32(1):268–274.

[118] Peng Y, Leung HCM, Yiu SM, et al. IDBA-UD: a *de novo* assembler for single-cell and metagenomic sequencing data with highly uneven depth. Bioinformatics. 2012;28(11):1420–1428.

[119] Dierckxsens N, Mardulyn P, Smits G. NOVOPlasty: *de novo* assembly of organelle genomes from whole genome data. Nucleic Acids Res. 2016;gkw955.

[120] Donath A, Jühling F, Al-Arab M, et al. Improved annotation of protein-coding genes boundaries in metazoan mitochondrial genomes. Nucleic Acids Res. 2019;47(20):10543–10552.

[121] Lang BF, Laforest M-J, Burger G. Mitochondrial introns: a critical view. Trends in Genetics. 2007;23(3):119–125.

[122] Chan PP, Lowe TM. tRNAscan-SE: Searching for tRNA Genes in Genomic Sequences. 2019. p. 1–14.

[123] Ontiveros-Palacios N, Cooke E, Nawrocki EP, et al. Rfam 15: RNA families database in 2025. Nucleic Acids Res. 2025;53(D1):D258–D267.

[124] Emms DM, Kelly S. OrthoFinder: phylogenetic orthology inference for comparative genomics. Genome Biol. 2019;20(1):238.

[125] Flynn JM, Hubley R, Goubert C, et al. RepeatModeler2 for automated genomic discovery of transposable element families. Proceedings of the National Academy of Sciences. 2020;117(17):9451–9457.

[126] Tarailo-Graovac M, Chen N. Using RepeatMasker to Identify Repetitive Elements in Genomic Sequences. Curr Protoc Bioinformatics. 2009;25(1).

[127] Stamatakis A. RAxML-VI-HPC: maximum likelihood-based phylogenetic analyses with thousands of taxa and mixed models. Bioinformatics. 2006;22(21):2688–2690.

[128] Kohany O, Gentles AJ, Hankus L, et al. Annotation, submission and screening of repetitive elements in Repbase: RepbaseSubmitter and Censor. BMC Bioinformatics. 2006;7(1):474.

[129] Rice P, Longden I, Bleasby A. EMBOSS: The European Molecular Biology Open Software Suite. Trends in Genetics. 2000;16(6):276–277.

[130] Danecek P, Bonfield JK, Liddle J, et al. Twelve years of SAMtools and BCFtools. Gigascience. 2021;10(2).

[131] Buchfink B, Reuter K, Drost H-G. Sensitive protein alignments at tree-of-life scale using DIAMOND. Nat Methods. 2021;18(4):366–368.

