## Supplementary figures and images for "The genome assembly of *Sphaerospora molnari* provides novel insights into the rapid evolution and diversification of unique lineage-specific gene groups in myxozoan parasites"

### Supplementary Figure 1

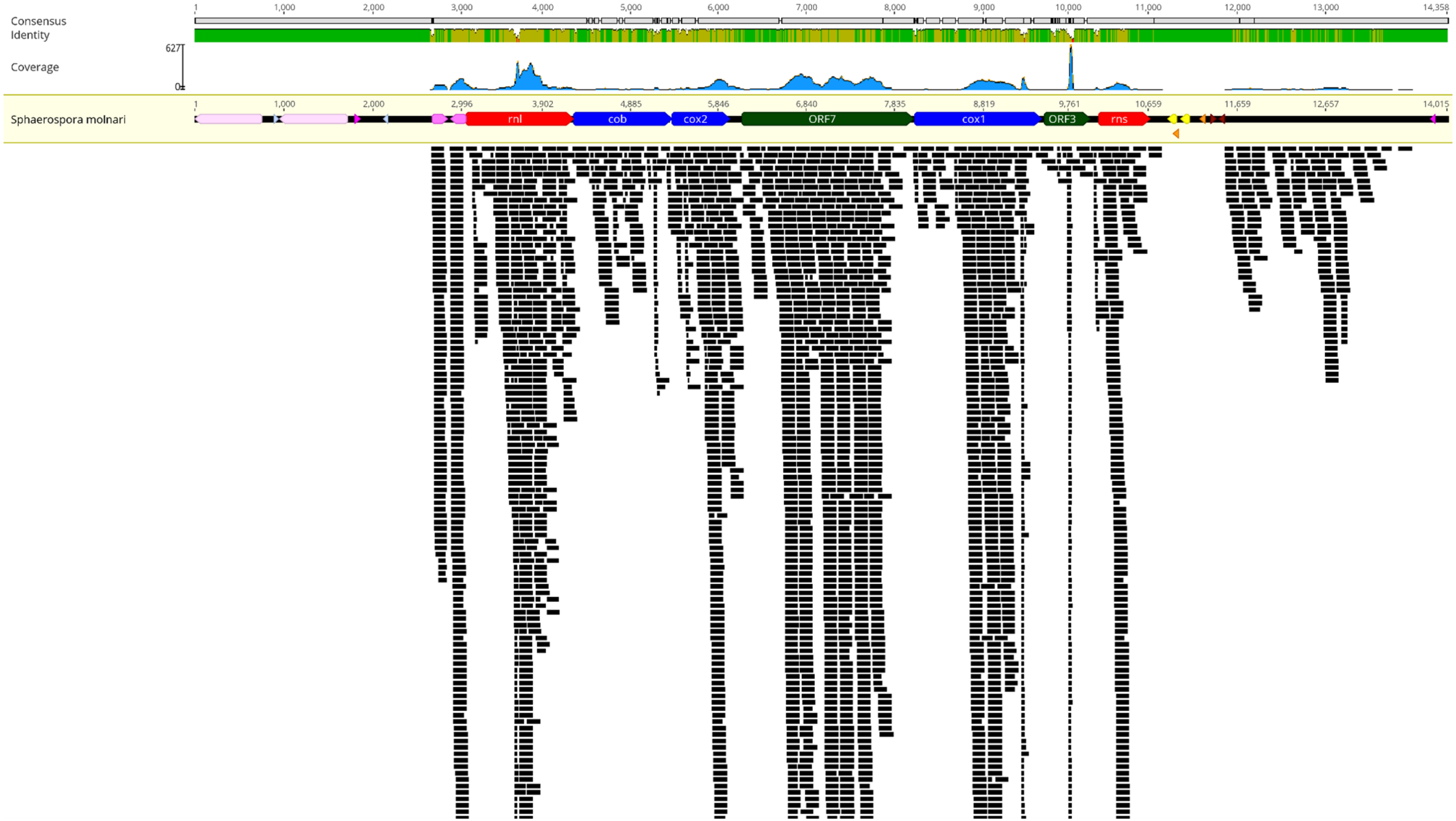

### Supplementary Figure 2

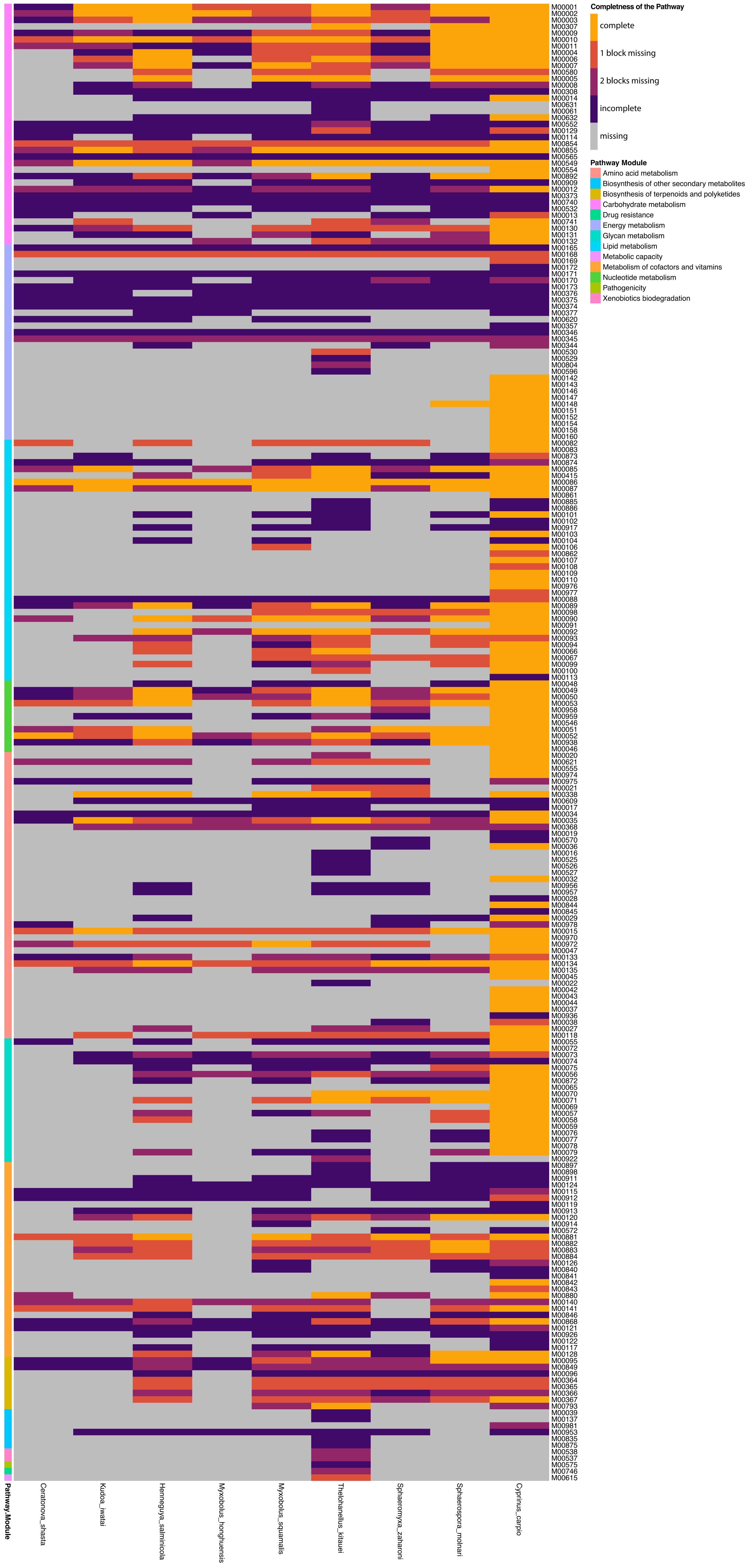

### Supplementary Figure 3

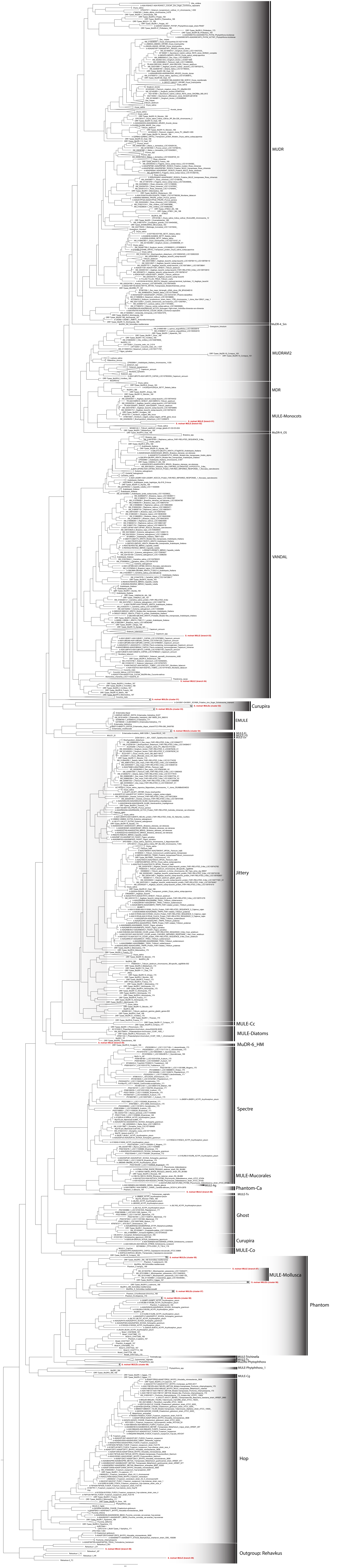
